# Cell-Graph Compass: Modeling Single Cells with Graph Structure Foundation Model

**DOI:** 10.1101/2024.06.04.597354

**Authors:** Chen Fang, Zhilong Hu, Shaole Chang, Qingqing Long, Wentao Cui, Wenhao Liu, Cong Li, Yana Liu, Pengfei Wang, Zhen Meng, Jia Pan, Yuanchun Zhou, Guihai Feng, Linghui Chen, Xin Li

## Abstract

Inspired by the advancements in pre-trained Large Language Models, there has been a surge of studies in the Life Sciences focusing on constructing foundation models with large scale single-cell RNA-seq data. These studies typically involve pre-training a transformer model on large-scale single-cell sequencing data, followed by fine-tuning for a variety of downstream tasks, achieving notable performance. However, these models all share a common short-coming: to utilize the transformer architecture, originally designed for textual data, they artificially impose a sequential structure on genes within cells, simplifying the complex interactions between genes. Furthermore, they focus solely on transcriptomic data, neglecting other relevant biological information. To address these issues, here we introduce Cell-Graph Compass (CGC), the first foundational model that leverages graph structures to model single cells and describes cells from multiple perspectives, including transcriptional profiles, gene text summaries, transcription factor regulatory networks, gene co-expression patterns, and gene positional relationships. By incorporating self-attention mechanisms, we pretrained the model on 50 million human single-cell sequencing data, resulting in a robust digital representation of cells. Extensive downstream experiments demonstrate that our approach can capture meaningful biological knowledge and achieve superior results in various problem scenarios, achieving the state-of-the-art (SOTA).

## 1 Introduction

Investigating the regulatory mechanisms between genes enhances our understanding of biological processes and plays a pivotal role in interpreting cellular functions, treating diseases, and promoting biotechnological innovations. Due to the intricate nature of biological networks, compounded by the prohibitive cost of wet-lab experiments, researchers urgently require efficacious computational simulation methods to help decipher gene regulatory mechanisms. Deep learning, at the forefront of Artificial Intelligence, has emerged as a research hotspot in recent years. However, its data-driven essence often constrains its utility in scenarios with limited data. The ”pre-train & fine-tune” paradigm,^1–5^originating from Nature Language Processing (NLP), offers a compelling solution. This kind of approaches generally involve pre-training a foundational model on extensive unlabeled datasets, followed by fine-tuning on small, task-specific datasets, thereby transferring generalized insights to domain-specific expertise. In parallel, the advancements in single-cell sequencing technologies and associated research^6–8^have amassed a wealth of single-cell transcriptomic data, providing a robust data resource for training foundational models in this domain.

Recently, pioneering efforts have been made in this arena. Geneformer^9^ introduced the concept of transfer learning into the single-cell domain for the first time. scGPT^10^ adopted generative pre-training to develop a universal foundational model. scFoundation^11^ creatively devised a read-depth-aware pre-training task to mitigate the impact of data quality… These progress represents significant breakthroughs in foundational models within the biological field. However, a common shortcoming exists in current models: an unwavering reliance on the transformer architecture,^1^ which was primarily designed for handling textual sequence data within the NLP domain. To meet the input demands of transformers, cells are conceptualized as “sentences” and genes as “words”. Nevertheless, unlike text, there is no inherent sequential structure among genes, and representing them in a serialized manner is a limited modeling approach. For such multidimensional relationships, modeling cells as graph structures may be more appropriate and can more easily represent the complex interrelationships in natural multiscale biological processes. Moreover, previous works have focused exclusively on extracting information from transcriptional expression profiles, neglecting the wealth of biological prior knowledge and previous research findings, which makes their models overly rely on the quality of collected sequencing data. The incorporation of biological prior knowledge may not only yield more robust results but also help direct the model to accurately capture and represent biological mechanisms.

In this paper, we introduce Cell-Graph Compass (CGC), a graph-based, knowledge-guided foundational model with large scale single-cell sequencing data. CGC conceptualizes each cell as a graph, with nodes representing the genes it contains and edges denoting the relationships between them. We employ a Graph Neural Network (GNN) architecture^12–14^ that integrates six different types of features and utilizes the message-passing mechanisms along with self-attention mechanisms to jointly learn the embedding representations of all genes. Our model is pre-trained on fifty million human single-cell sequencing data from ScCompass-h50M and validate its effectiveness across a broad range of downstream experiments. We conducted gene identity recognition and gene regulatory network inference tasks under a zero-shot setting, validating that CGC captured biologically meaningful information during pretraining. We fine-tuned CGC for various downstream tasks, including batch effect correction, cell type annotation, single-cell gene perturbation, and bulk gene knockout prediction. CGC consistently exhibited exceptional performance across all applications, underscoring the efficacy of constructing a pretrained foundational model by integrating graphs with biological knowledge.

## 2 Results

### 2.1 The Graph-based foundation model overview

CGC combines the structure of Graph Neural Networks (GNN) with the transformer architecture, modeling cells in an intuitive manner while incorporating various biological information. The architecture of our model consists of four interconnected modules, as depicted in Fig.1(a): an Encoder module, a GNN module, a Transformer module, and a Decoder module. In this framework, the Encoder module is responsible for encoding and integrating various feature data into the nodes’ and edges’ representations. The GNN module employs a message-passing mechanism to propagate genes’ information across connected nodes while the Transformer module leverages its self-attention mechanism to comprehensively learn and refine the genes’ feature representations, exploring the complex interactions and relationships among genes. In the final stage, tailored to suit different application needs, distinct Decoder modules have been designed to convert the hidden vectors outputted by the Transformer module into the anticipated results, thus unlocking the model’s vast potential across a range of biological tasks.

**Fig 1:**
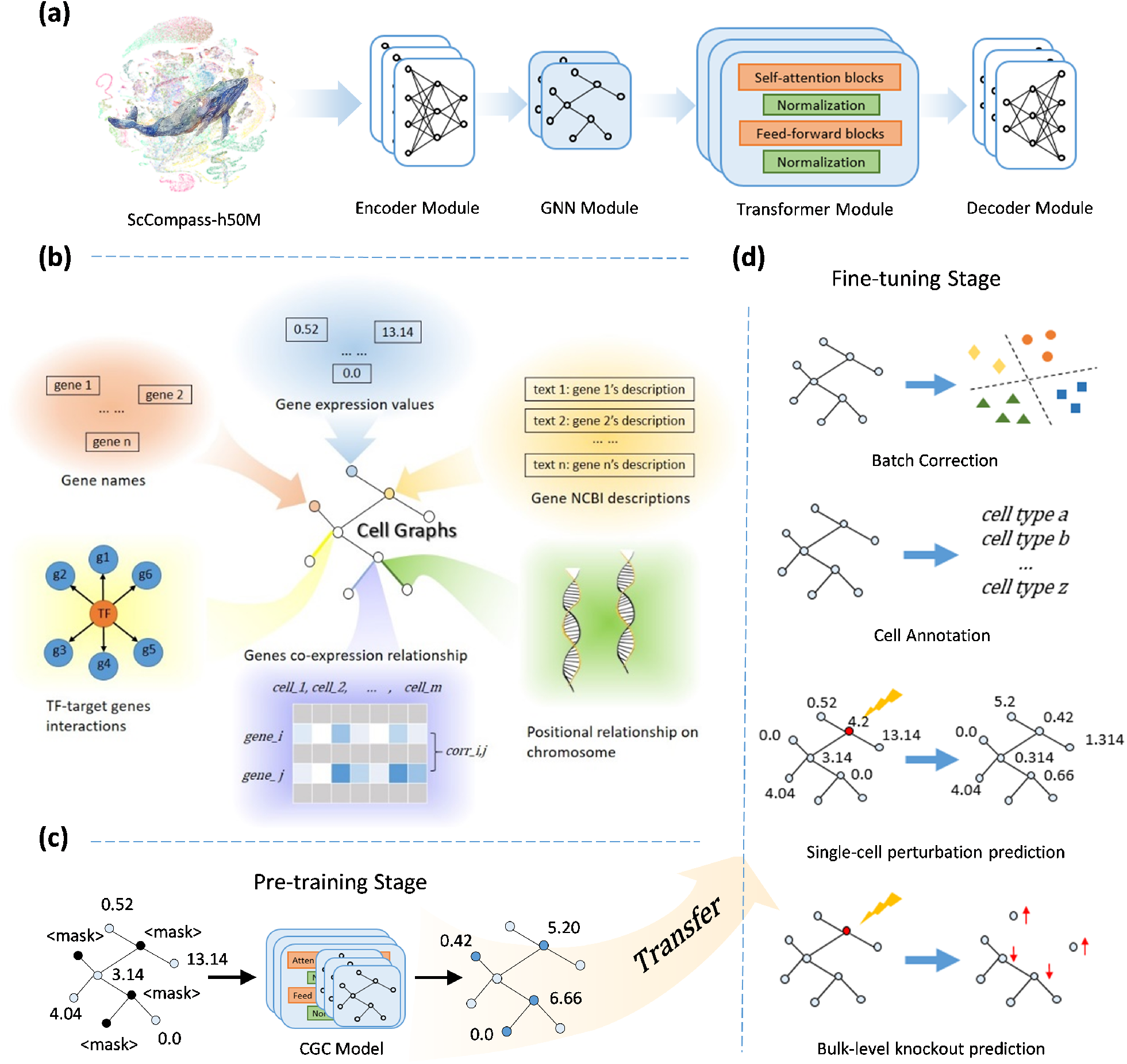
Overview of Cell-Graph Compass (CGC) methodology. (a) Model Architecture: The CGC model consists of four components: an Encoder, a Graph Neural Network (GNN), a Transformer, and a Decoder module. (b) The construction of cell graphs: CGC utilizes gene tokens, gene transcription expression values, and gene text descriptions as node features and constructs edges based on transcription factor-target gene Interactions, gene co-expression relationships, and genes’ positional relationship on chromosome, with the GNN module to synthesize and vectorize these features. (c) Pre-training stage: CGC learns general knowledge about genes through masked training on fifty million human single-cell sequencing data. (d) Fine-tuning stage: CGC reaches state-of-the-art outcomes in a plethora of downstream tasks, exemplified by four instances in this paper: batch effect correction, cell type annotation, single-cell gene perturbation prediction, and general bulk gene knockout prediction.

In cell biology, the transcriptional expression of genes within a single cell serves as a snap-shot of that cell’s state and function. This concept forms the fundamental basis of recent foundation model studies,^9–11,15, 16^ which generate embedding representations of genes and cells based on single-cell RNA-seq data. Beyond this, CGC utilizes scBERT,^15^ a specialized language model from the biomedical field, to digitize textual descriptions of genes from the NCBI database.^8^ These digitized descriptions serve as one-dimensional features for the nodes, thereby providing the model with summarized gene information from previous research. As a result, we incorporate a total of three distinct node features: gene tokens, gene transcriptional expressions and gene text descriptions. To construct the edges of cell graphs, we collected information on gene interrelations or interactions from three different dimensions: previously summarized transcription factor-target gene interactions, statistically quantified gene co-expression patterns, and the positional information of genes on chromosome, as illustrated in Fig.1(b). These edges direct the message-passing in the GNN module, which employs an iterative co-learning process for all nodes and edges (see Methods 4.2), exemplifying a method of local feature extraction. Subsequently, the Transformer module considers the attention weights of all gene pairs from a global perspective. This combined approach of considering both local and global connections not only focuses on gene relations emphasized by the edges but also aids in uncovering potential unknown relationships.

For pre-training, we utilized approximately 50 million human single-cell transcriptomes from ScCompass-h50M and each entry was subjected to strict quality control and normalized procedures. Our pre-training strategy, as shown in Fig.1(c), randomly masks 40% of gene expression values for each input and employs the remaining 60% to infer overall cellular state and to predict the expression of the masked genes. Additionally, a global node representing the entire cell was introduced, connected to all gene-specific nodes, thereby explicitly modeling the cellular state. After pre-training, CGC could be fine-tuned for abundant downstream tasks, as illustrated in Fig.1(d). Its flexible modular design allows for simple adjustments to accommodate different application scenarios, making it highly adaptable for fine-tuning to address specific problems.

### 2.2 Through pre-training, CGC acquires biologically meaningful knowledge

Before fine-tuning the CGC model for specific downstream tasks, it is necessary to ensure through analysis that the pre-trained model has achieved the anticipated effects. Specifically, a successful pre-trained model should fundamentally understand genes and their interactions. Therefore, in this session, we investigated whether CGC can recognize special genes and effectively infer Gene Regulatory Networks (GRNs) by exploring the hidden layer space of the pre-trained model.

In the CGC’s Encoder module, we use gene tokens to represent all the genes encountered during pre-training and obtain a general embedding for each token through this process. These embeddings reflect the model’s memory of these genes from the pre-training stage, forming the basis for our subsequent experiments.

To test whether the pre-trained model can distinguish specific genes, we selected six gene classification datasets covering various aspects such as gene dosage sensitivity,^17–19^chromatin dynamics,^20, 21^ and network dynamics^22, 23^ (details in Methods 4.4). We also compared three other methods for generating general gene representations: Gene2vec,^24^ which creates embeddings based on gene co-expression patterns; BioBERT,^25^ a representative language model for the biomedical field, which generates embeddings from gene descriptions in the NCBI database^8^ ; and randomly generated gene embeddings. We then used a random forest^26, 27^ classifier to perform gene classification based on the embeddings produced by each of these four methods. Results of the five-fold cross-validation experiments showed, as illustrated by the Area Under Curve (AUC) values in Fig.2(a) and the Receiver Operating Characteristic (ROC) curves in Supplementary Fig.S1(a)-(f), that randomly generated embeddings had AUC values around 0.5, equivalent to random guessing, while the results of the other three methods were significantly higher than 0.5, confirming their embeddings indeed captured gene-related information. Notably, CGC achieved the best performance across all six datasets, validating our pre-training on fifty million cell graphs enabled CGC to gain a certain level of understanding of genes.

**Fig 2:**
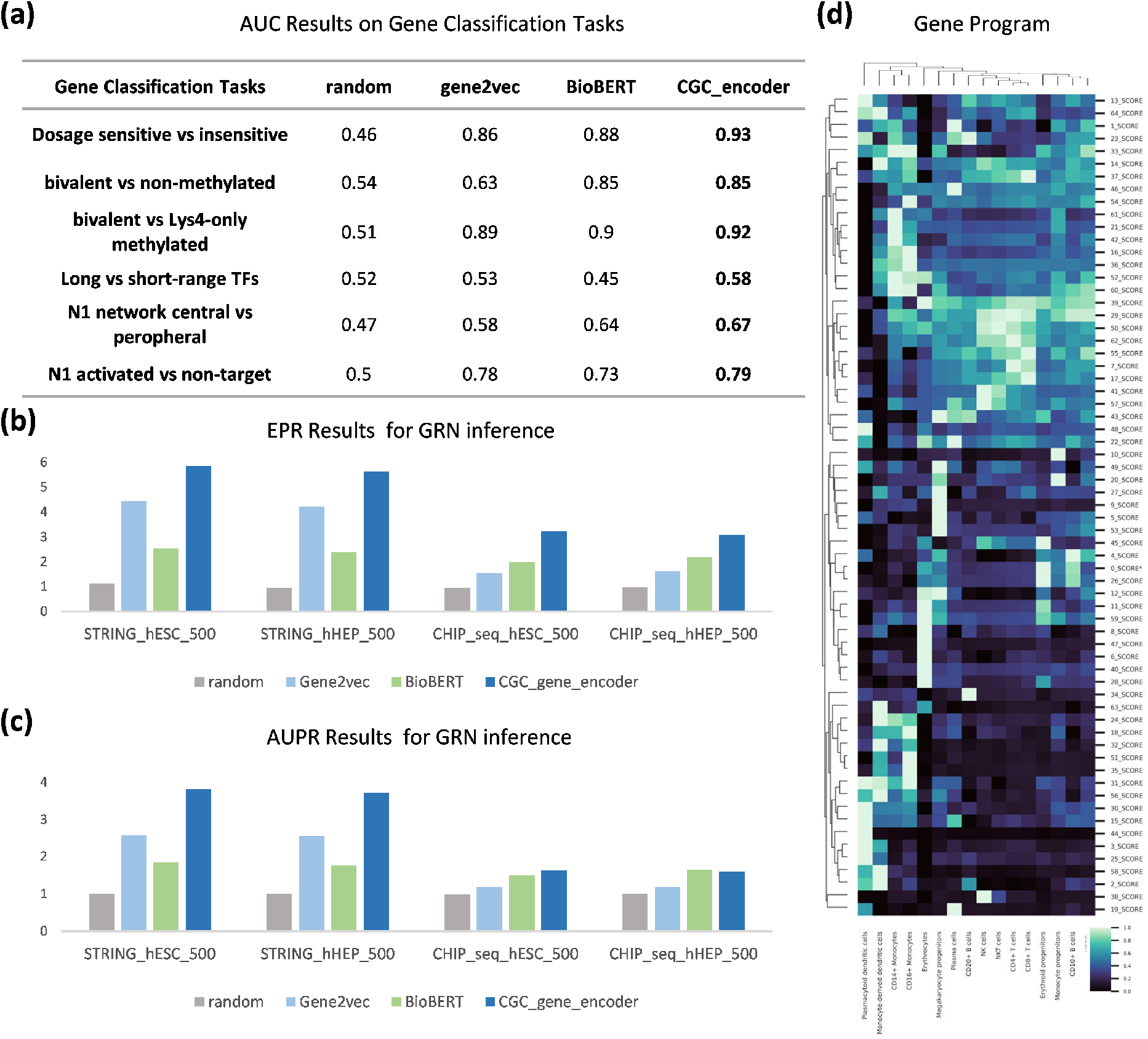
Pre-training endows CGC with biologically meaningful knowledge. By dissecting the latent space of the pre-trained model, we obtain a universal embedding representation for each gene. We evaluate CGC’s ability to capture biological knowledge by assessing whether these embeddings can pinpoint the identity of specific genes and if they encompass information about Gene Regulatory Networks (GRNs). (a) Results of six gene classification experiments. We selected two cutting-edge methods for extracting universal gene embeddings along with randomly generated embeddings as baselines. (b) Results of GRN inference. The Early Precision Ratio (EPR) metric measures the overlap between the model-reconstructed GRN and the ground truth. (c) The Area Under the Precision–Recall Curve (AUPRC) ratio metric evaluates the extent to which the model predicts the ground truth GRN. (d) The selective activation of the gene program extracted by CGC across different cell types.

Next, we conducted GRN inference experiments using data from STRING^28^ and ChIP-seq^29–31^ databases, covering four cell types. Inspired by Pratapa et al.,^32^ we selected 500 and 1000 transcription factors (TFs), respectively, creating 8 datasets in total. Following the test approach of DeepSEM,^33^ We used the cosine similarity of gene pair embeddings to judge their potential regulatory relationships. We evaluated the quality of GRNs generated by these four methods using EPR and AUPR metrics (detailed in Methods 4.4). The results, as shown in Fig.2(b) & (c) along with Supplementary Fig.S1(g) & (f), indicate that across all 8 datasets, the embeddings provided by CGC consistently reconstructed the most complete GRNs, demonstrating our pre-training enabled CGC to learn about the interactions between genes to some extent.

Moreover, we noticed that in gene classification and GRN inference experiments, embeddings generated by BioBERT performed comparably to CGC’s gene token embeddings across several datasets, suggesting that embeddings generated by language models based on text descriptions can also provide useful biological information. This supports our intention of using them as one input feature for nodes.

CGC can also be used to extract meaningful gene programs. We clustered genes into several programs based on gene token embeddings and analyzed their selective expression in the human immune tissues dataset.^34^ As shown in Fig.2(d) and Supplementary Fig.S2, functionally similar genes were grouped into the same program, with clear expression differences between different programs. For example, gene program 2 extracted by CGC included the CD74 gene and a group of HLA II class genes, which are highly expressed in Monocyte-derived dendritic cells. In fact, CD74, HLA II class genes, and dendritic cells are closely linked,^35^ crucial for initiating and regulating specific immune responses. Similarly, gene program 12 contained a group of histone-related genes, playing a key regulatory role in the development of red cells and megakaryocytes^36^ and the maturation of their precursor cells. These findings demonstrate that gene token embeddings provided by CGC can reveal functional gene clusters, further evidencing that CGC’s pre-training can learn biologically meaningful information of gene features and programs..

### 2.3 Fine-tuning the pre-trained CGC model to mitigate batch effects

CGC integrates single-cell sequencing data from various datasets, and thus, the first challenge we encounter is the batch effect^37^ which refers to the technical noise arising due to differences in sequencing platforms, experimental instruments, or analysis methods, which might obscure the biological signals of interest. Correcting batch effects to acquire useful biological knowledge is a fundamental requirement for foundational models in the single-cell domain.

Cell clustering is a common downstream task that reflects models’ ability to correct for batch effects. The objective of this task is to derive high-dimensional features that effectively reflect cell type information while minimizing the impact of batch variations as much as possible. To evaluate the capability of CGC on batch correction task, we conducted validations across three publicly accessible datasets: the Perirhinal Cortex (P.C.) dataset,^38^ containing data from ten cell types across two batches; the PBMC dataset^39^ with two batches and nine cell types; and the COVID-19 dataset,^40^ comprising 18 batches and 39 cell types. We fine-tuned CGC on these datasets using the same self-supervised masked training approach as in the pre-training stage, aiming to transfer the general knowledge acquired from ScCompass-h50M to specific downstream tasks. We employed four metrics for evaluating clustering performance: NMI, ARI, ASW, and GraphConn, and used their average as the overall score. The first two metrics assess clustering by cell types (reflecting the model’s capacity to remain biological signal), while the latter two measure the mixing of data from different batches (reflecting the ability to mitigate batch variations). Detailed definitions of these metrics can be found in Methods 4.4. We chose scVI,^41^ scFoundation,^11^ and scGPT^10^ as baselines. scVI is a variational autoencoder model specifically for single-cell transcriptome data analysis, while scFoundation and scGPT are recently introduced large foundation models based on the “pretraining & fine-tuning” paradigm. Fig.3 (a)(e), and Supplementary Fig.S3(c) display UMAP plots of cell embeddings generated by each model on these three datasets respectively. All models could broadly distinguish between cell types compared to the original input, with CGC forming more compact clusters and clearer separations between them. Fig.3 (b)(f), and Supplementary Fig.S3(d) show the specific quantitative outcomes for each model. CGC achieved the best scores across both small-scale and large datasets like COVID-19, affirming its ability to counteract batch effects and its universality. Notably, while scFoundation excelled in the COVID-19 dataset, its performance on the P.C. dataset was subpar, possibly due to its pre-training dataset being more aligned with the COVID-19 dataset’s distribution than with P.C.’s. In contrast, CGC consistently yielded good results across all three datasets, which may benefit from our comprehensive pre-training dataset and leveraging numerous biological priors to address potential biases.

To validate the effectiveness of our graph-based approach, we conducted ablation experiments on CGC’s graph structure and pre-training process. Fig.3 (c)(g) and Supplementary Fig.S3(e) show the outcomes of the graph structure ablation on three datasets, respectively, with a notable decline in performance across all metrics after removing the graph structure and GNN module. This confirms the effectiveness of modeling genes and cells through graph structures. Fig.3 (d)(h) and Supplementary Fig.S3(f) present the results of the pre-training process ablation, demonstrating significantly poorer performance for models trained from scratch compared to those pre-trained. Given CGC’s use of a GNN module to integrate various biological features, without pre-training, the model may struggle to synthesize and utilize such diverse information.

**Fig 3:**
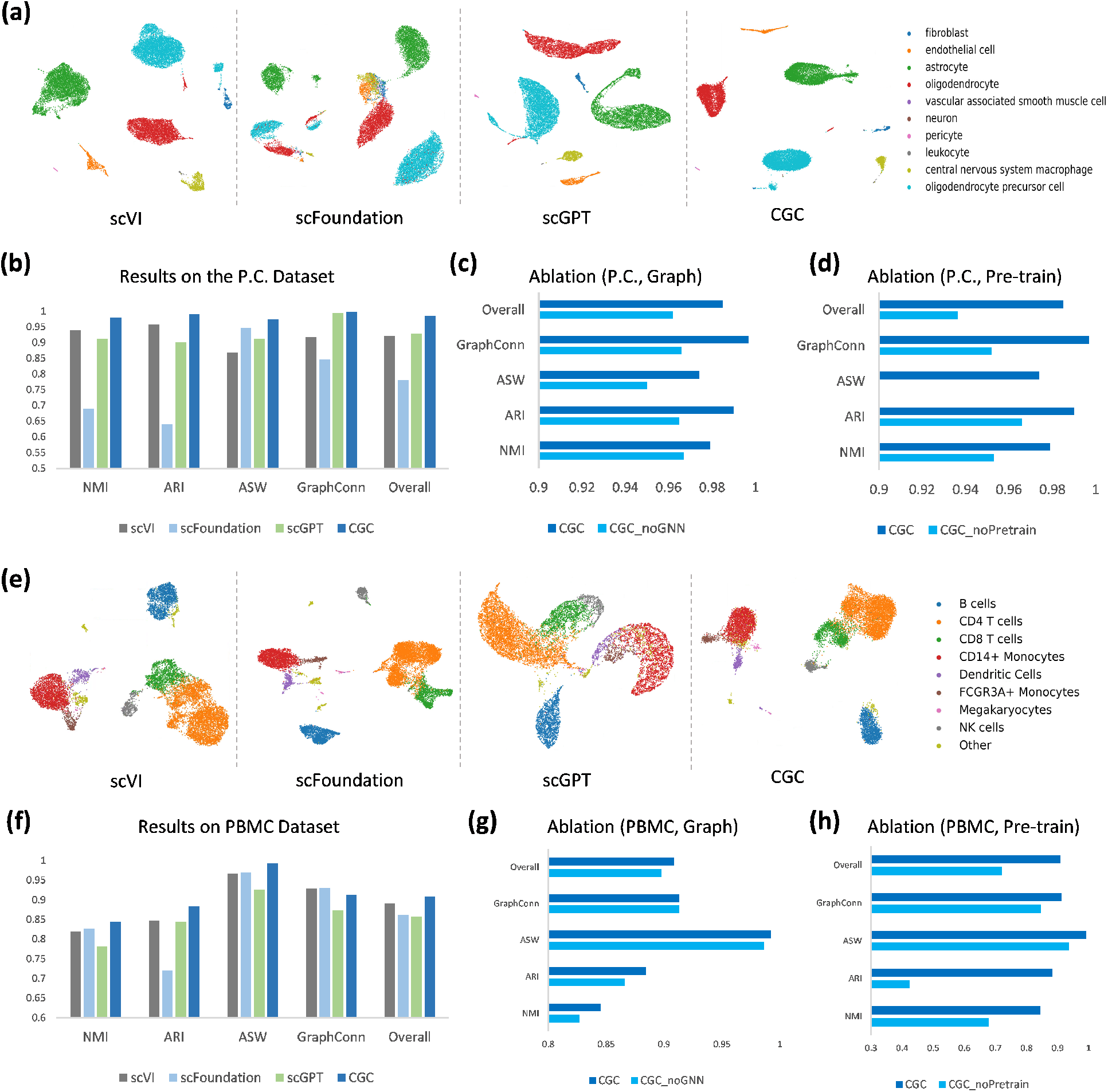
Finetuned CGC diminishes the impact of batch effects. On the PCortex dataset, (a) UMAP plots of cell embeddings generated by CGC and the baseline models, colored by the ground truth cell types. (b) Benchmark results for CGC and the baseline models. (c) Ablation study on the graph structure of CGC. (d) Ablation study on the pre-training process of CGC. On the PBMC dataset, (e) UMAP plots of cell embeddings generated by CGC and the baseline models. (f) Benchmark results for CGC and the baseline models. (g) Ablation study on the graph structure of CGC. (h) Ablation study on the pre-training process of CGC.

### 2.4 CGC exhibits excellent performance in cell type annotation

In this section, we continue to explore the performance of CGC in downstream tasks at the cellular level. Cell type annotation, as a pivotal part of single-cell data analysis, plays an essential role in understanding the diverse functionalities and characteristics of cells. While numerous advanced annotation methods^15,42, 43^ have been proposed and achieved notable results, they generally rely solely on single-cell transcriptome data, leading to a limited understanding of cellular complexity. CGC distinguishes itself by both learning the cell’s context through intracellular gene expression profiles and integrating various biological priors at the same time, which makes it one of the most comprehensive and effective cell annotation methods among state-of-the-art technology.

To get an embedding that can comprehensively represent the whole cell state, we have additionally designed a global node. This node is connected to all other nodes representing genes within the graph, and the connection weight parameters are automatically learned during the training process. In the fine-tuning phase, we used an Multi-layer Perceptron (MLP) as the classification head to output the predicted cell type based on this cell embedding.

We validated our model on three publicly available datasets, each containing a reference set and a query set with no overlap between them. We fine-tuned the CGC model on the reference sets and tested it on the query sets, using Accuracy, Precision, Recall, and F1 score as metrics. We selected scANVI,^44^ Geneformer,^9^ and scGPT^10^ as baselines. Our experiments began with the Multiple Sclerosis (M.S.)^45^ dataset, which includes 17 cell types with the reference set derived from healthy human immune cells and the query set from M.S. patients. Fig.4(b) shows the distribution of the M.S. query set and CGC’s prediction results, which closely match the true labels qualitatively. Fig.4(c) displays the quantitative test results of CGC compared to baseline models, indicating that pre-trained foundation models (scGPT, Geneformer, CGC) outperform the task-specific model trained from scratch (scANVI), validating the effectiveness of the pre-training strategy. Moreover, the CGC model, based on cell graphs, significantly outperforms Transformer-only models (scGPT, Geneformer), highlighting the beneficial effect of the GNN architecture on single-cell foundation models. Fig.4(d) details CGC’s classification results for each cell type, showing normalized results on the left and absolute numbers on the right. Except for categories with extremely few samples (such as endothelial cells) where CGC fails to predict, CGC provides over 90% accuracy for the vast majority of categories.

In more in-depth testing, we first conducted ablation experiments on the graph structure and the GNN module. As shown in Fig.4(e), the addition of the GNN module to the Transformer module showed superior performance compared to the Transformer-only model. Furthermore, we conducted ablation experiments on the CGC pre-training process. When the model was trained from scratch, a significant decrease in annotation effectiveness occurred, as shown in Fig.4(f), highlighting the importance of pre-training and demonstrating that the graph-based model greatly benefits from pre-training data for optimal performance.

**Fig 4:**
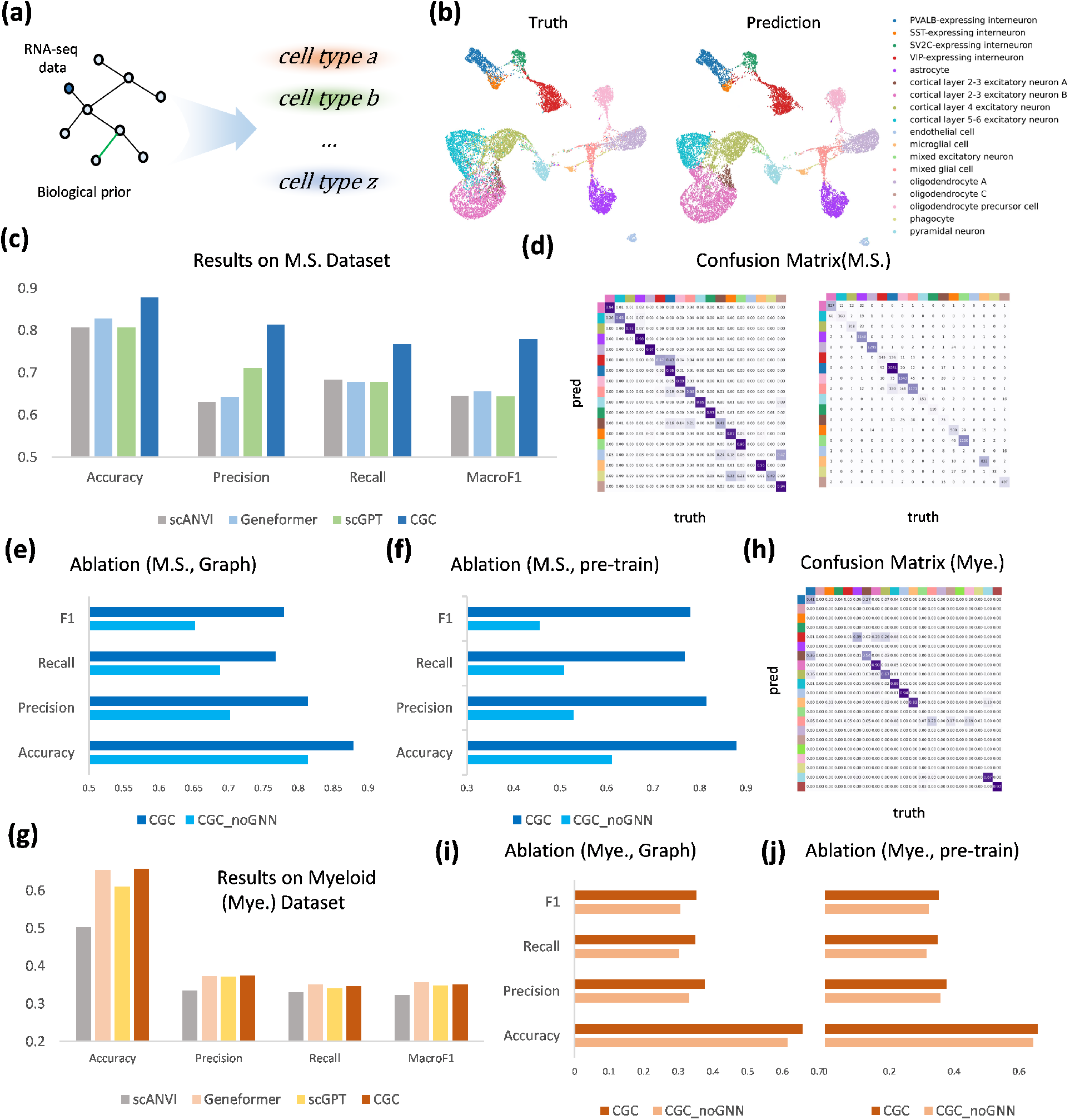
CGC boosts cell type annotation. (a) Schematic of the cell type annotation task. On the Multiple Sclerosis (M.S.) dataset, (b) UMAP visualization of cell embeddings generated by CGC, colored by ground truth cell types (left) and CGC prediction results (right). (c) Quantitative evaluation of cell annotation by CGC and baseline models. (d) Confusion matrix between cell types predicted by CGC and ground truth labels, with normalization on the left and absolute values on the right. (e) Ablation study on the graph structure of CGC. (f) Ablation study on the pre-training process of CGC. On the Myeloid (Mye.) dataset, (g) Quantitative evaluation of cell annotation by CGC and baseline models. (h) Confusion matrix between cell types predicted by CGC and ground truth labels. (i) Ablation study on the graph structure of CGC. (j) Ablation study on the pre-training process of CGC.

Next, we conducted experiments on the Myeloid (Mye.)^46^ and hPancreas^42^ datasets. In the Mye. dataset, the reference set originated from six types of cancer, including 21 cell types; the query set came from three other types of cancer, including 11 cell types. In the hPancreas dataset, the reference set came from two small datasets^47, 48^ of human pancreas cells, including 13 cell types; the query set came from three other small datasets,^49–51^ including 11 cell types, one of which was not seen in the reference set. The results, shown in Fig.4(g) to (j) and Supplementary Fig.S4, indicate that CGC still achieved the best annotation results. The ablation experiments on both the graph structure and the pre-training process also had significant effects. Additionally, Geneformer showed outstanding performance in the Mye. dataset but significantly lagged in the hPancreas dataset. scANVI, as a small model trained from scratch, although generally lagging behind pre-trained foundation models in terms of overall annotation performance, still surpassed Geneformer and scGPT in several evaluation metrics, revealing that neither pre-trained models nor task-specific models have the ability to perform well in all scenarios. In contrast, CGC, with its multitude of biological priors, compensates for the distribution differences between pre-training data and downstream task datasets, offering reliable performance and scalability.

### 2.5 CGC excels at predicting single-cell gene perturbation response

After exploring the downstream tasks at the cellular level, we proceed to examine the performance of CGC in gene-level application scenarios. Precise prediction of gene perturbation responses is crucial for unveiling cellular biological traits, understanding disease mechanisms, and drug development, among other fields.^52–54^ The advancement of gene editing^55^ technologies and Perturb-seq^56^ has laid the groundwork for the application of deep learning method.^57–59^ GEARS^60^ is one of the most advanced perturbation response prediction models to date. Similar to our concept, it also adopts a GNN architecture and utilizes gene co-expression relationships and gene ontology knowledge to construct graphs, aim for predicting the perturbation response of genes. In contrast, CGC utilizes a broader and more complex array of biological features for graph construction and undergoes pre-training on fifty million cell graphs to acquire a general understanding of genes’ relationship in advance.

Our empirical evaluation utilized a public dataset named Norman,^61^ which includes 105 single-gene and 131 double-genes perturbations. We adopted the same dataset split method as GEARS, ensuring that the test set contains perturbation conditions (which gene/s are perturbed) not present in the training set. To evaluate model performance, we chose GEARS and scGPT as baselines. We used the mean squared error (mse) and pearson correlation coefficient (corr) of gene expression values after perturbation, along with the correlation coefficient of the change in gene expression (corr delta), as evaluation metrics and tested them across all genes and the top 20 differentially expressed (DE) genes separately. As shown in Fig.5 (a), the two transcriptomic foundation models have a significant advantage over GEARS, highlighting the importance of pre-training in this task. Additionally, CGC achieved a clear improvement over scGPT, further proving the advantage of the graph structure foundation model. Fig.5(b) offers a clearer visualization of CGC’s enhancement, where each point corresponds to a gene in the test set. Distinct colors differentiate the predictions made by various models, and lines are drawn to fit the points corresponding to each model, illustrating the comparative performance visually. Furthermore, we also examined the three models’ ability to predict the direction of gene expression change after perturbation (up, unchanged, down), as shown in Fig.5(c), CGC consistently produced the best results when evaluating within different numbers of DE genes.

When conducting gene editing experiments, the same perturbation condition usually produces a number of sequencing samples. This is because, despite the high precision of gene editing technologies like CRISPR-Cas9,^55^ there is still some variability and uncertainty in this process. Therefore, to ensure the reliability of the results, experimenters tend to edit a large amount of cells under the same perturbation condition at the same time.

Corresponding to this situation, we revisited the predictive ability of all models from a different perspective. We treat the entire set of sequencing samples under each perturbation condition as a single distribution and calculate the average for each distribution, encompassing both actual and predicted values. Then, we tested these models’ predictions for these distributions using the same metrics. This testing method overlooks the heterogeneity between cells but reduces the impact of uncertainty on the wet-lab experiment, focusing more on common results. Under this new testing perspective, as shown in Fig.5(e), scGPT’s performance declined, slightly below GEARS. This is because both CGC and scGPT model each sequencing sample, while GEARS directly models each distribution, thus having an advantage in this type of test. However, CGC still achieved the best results and gained a 3-point improvement over GEARS. This reaffirms the efficacy of CGC, highlighting its resilience to random fluctuations and the robustness and reliability of its predictive outcomes. Further, we conducted ablation experiments on CGC’s graph structure and pre-training process, with Supplementary Fig.S5 showing the results, as expected.

Additionally, we analyzed on the gene-by-gene basis of predictions. Supplementary Fig.S6 presents four instances, corresponding to the prediction results of all models under four specific perturbation conditions. In the diagram, ”boxes” represent the distributions of the true variation values of the top 20 DE genes across all samples under that perturbation condition, and the black dots represent the average values of these distributions. Different colored dots represent the average of all samples’ predicted values by different models under that perturbation condition. It is observable that CGC’s predictions are closest to the real average values in the vast majority of cases. We also summarized the probability of each model’s predictive points falling within the center of the ”boxes”, as shown in Fig.5(f), with CGC consistently outperforming baseline models as the center “range” gradually increases. In summary, CGC can achieve accurate and robust perturbation predictions, thanks to its graph-based pre-training.

**Fig 5:**
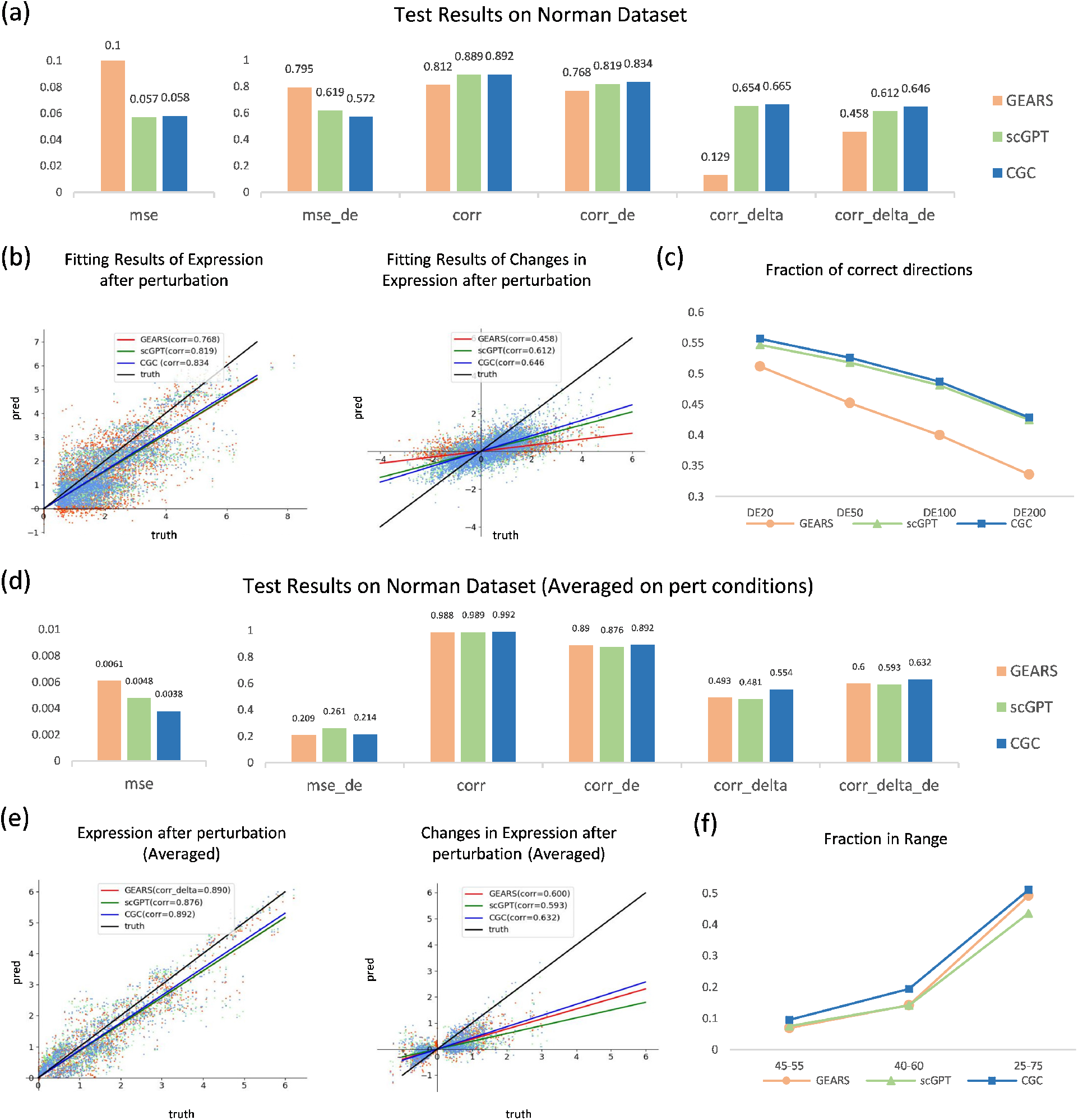
CGC accurately predicts single-cell gene perturbation responses. (a) Quantitative comparison of the perturbation response predictions between CGC and baseline models. (b) Scatter plot of all models’ predictions on post-perturbation gene expression values (left) and change values (right). The x-axis represents the true values, and the y-axis represents the predicted values, with each point corresponding to a gene in a sample from the test set, showcasing 1% of total genes. Lines of different colors fit the scatter plots of different models. (c) The correct prediction ratio of all models on the direction of gene change (increase, no change, decrease), with the x-axis representing the different number of DE genes included in the evaluation. (d) Quantitative comparison of the predictions between all models on the average response of genes under different perturbation conditions. (e) Scatter plot of all models’ predictions on the average post-perturbation gene expression values (left) and change values (right) by perturbation condition. Each point represents the average expression value of a gene under one type of perturbation condition. (f) The proportion of prediction points from all models that fall within the true distribution center for post-perturbation change amounts. The x-axis indicates the range of the “center” of the true distribution.

### 2.6 CGC innovatively explores the construction of a “universal” gene knockout prediction model

Single-cell sequencing technology provides detailed analysis of gene expression in individual cells, aiding research on cellular heterogeneity. However, due to its high cost, existing single-cell perturbation experiments^56,61–64^are generally limited to specific cell lines and a few perturbation conditions. Consequently, the perturbation prediction models designed are often only applicable to fixed cell lines and specific cellular states, losing their predictive power on new batches of experimental data. Designing a universal perturbation prediction model remains an open challenge.

Bulk sequencing offers a faster, more cost-effective alternative. Conducting gene perturbation studies at the bulk level allows for coverage of a wider variety of cellular states and a broader range of perturbation conditions. Although this method overlooks differences between individual cells, it focuses more on the gene expression information of the bulk overall, potentially revealing more universal and general gene regulation patterns. To study bulk gene perturbation, we collected over 3,300 experimental instances on mouse gene knockouts from the web, covering more than 27,000 genes and over 1,100 perturbation conditions (see Fig.6(b)). Since the data come from different wet labs, each sample can be considered to be in a different cellular state, necessitating the inference of cellular state information based on pre-knockout gene expression profiles. Approaches like GEARS cannot be applied to this context, hence it models the ”distribution” of the perturbation situation. In this scenario, each distribution consists of only one sample.

Bulk sequencing results are an aggregate average of multiple cells. In theory, knowledge obtained from single-cell data should be easily extendable to the coarser bulk level. To facilitate smoother knowledge transfer, we employed a ”two-step transfer” pre-training strategy. We first pre-trained the model on 20 million mouse single-cell sequencing data, followed by a second masked training on 300,000 bulk sequencing data. Finally, we fine-tuned on the collected 3,300 bulk knockout data to obtain the final bulk gene knockout model. Using this strategy, we initially attempted to directly predict the expression values of all genes post-knockout, employing the same model architecture and testing method as in the previous section. Fig.6(c) shows the performance of CGC and scGPT on the test set, with CGC demonstrating superior performance, showcasing our single-cell pre-trained model’s transferability at the bulk level.

Further, we attempted to predict the direction of gene expression changes after knockout. In this context, if the expression of a gene increased more than twofold post-knockout, it was defined as an ”up-regulated gene”; if it decreased more than twofold, it was defined as a ”down-regulated gene”; those not meeting these criteria were considered ”unchanged”. Consequently, in the bulk knockout dataset, on average, only about 4% of genes per sample belonged to either the up-regulated or down-regulated categories. Such extreme data imbalance led the model to predict that all genes remain unchanged because doing so would still achieve a high accuracy rate of up to 96%.

To prevent the model from taking this ”lazy” approach, we made a series of related designs. First, we replaced the original regression decoder with two classifiers to directly model the direction of change. As shown in Fig.6(a), the first classifier predicts whether a gene changes, while the second predicts whether a changing gene is up-regulated or down-regulated. Thus, the problem of data imbalance is mainly concentrated in the first classification task. For the first classifier, we used focal loss^65^ as the training objective to increase the weight of changed genes and employed negative sampling to balance the number of positive and negative samples. For comparison, we also replaced the output layer of scGPT with our dual-classifier structure and used the same loss for training. We employed three-category testing metrics to evaluate model performance, with results shown in Fig.6(d). It’s evident that although CGC showed significant improvement over the baseline, the F1 score still did not reach the desired effect, indicating the difficulty of this task. To explore the potential application of the model in actual experiments, we visualized the confusion matrix of the test results in Fig.6(e). It can be seen that, while maintaining an overall accuracy rate of over 90%, nearly half of all up- and down-regulated genes were correctly predicted, with the other half predicted as unchanged, and a very small proportion had their direction of change incorrectly judged. In other words, although CGC cannot predict all up- and down-regulated genes, the likelihood of it providing opposite answers is very low, suggesting that CGC can narrow down the experimental exploration range and take an important step forward in the quest to build precise prediction models.

More intricately, we tried predicting the magnitude of gene changes and formalized this as a five-category problem, predicting whether a gene remains unchanged, up-regulated twofold, down-regulated twofold, up-regulated fivefold, or down-regulated fivefold. The five-category task presented a more challenging hurdle, but compared to the baseline model, CGC still achieved a considerable improvement in performance (see Fig.6(f)). As shown in Fig.6(g), genes up-regulated fivefold were predominantly judged as either up-regulated or unchanged. Similarly, genes up-regulated twofold were primarily categorized as either up-regulated twofold or unchanged. This pattern was consistent for down-regulated genes as well. This means that the probability of incorrectly judging the direction of gene changes was very low, revealing the feasibility of CGC aiding actual biological experiments.

**Fig 6:**
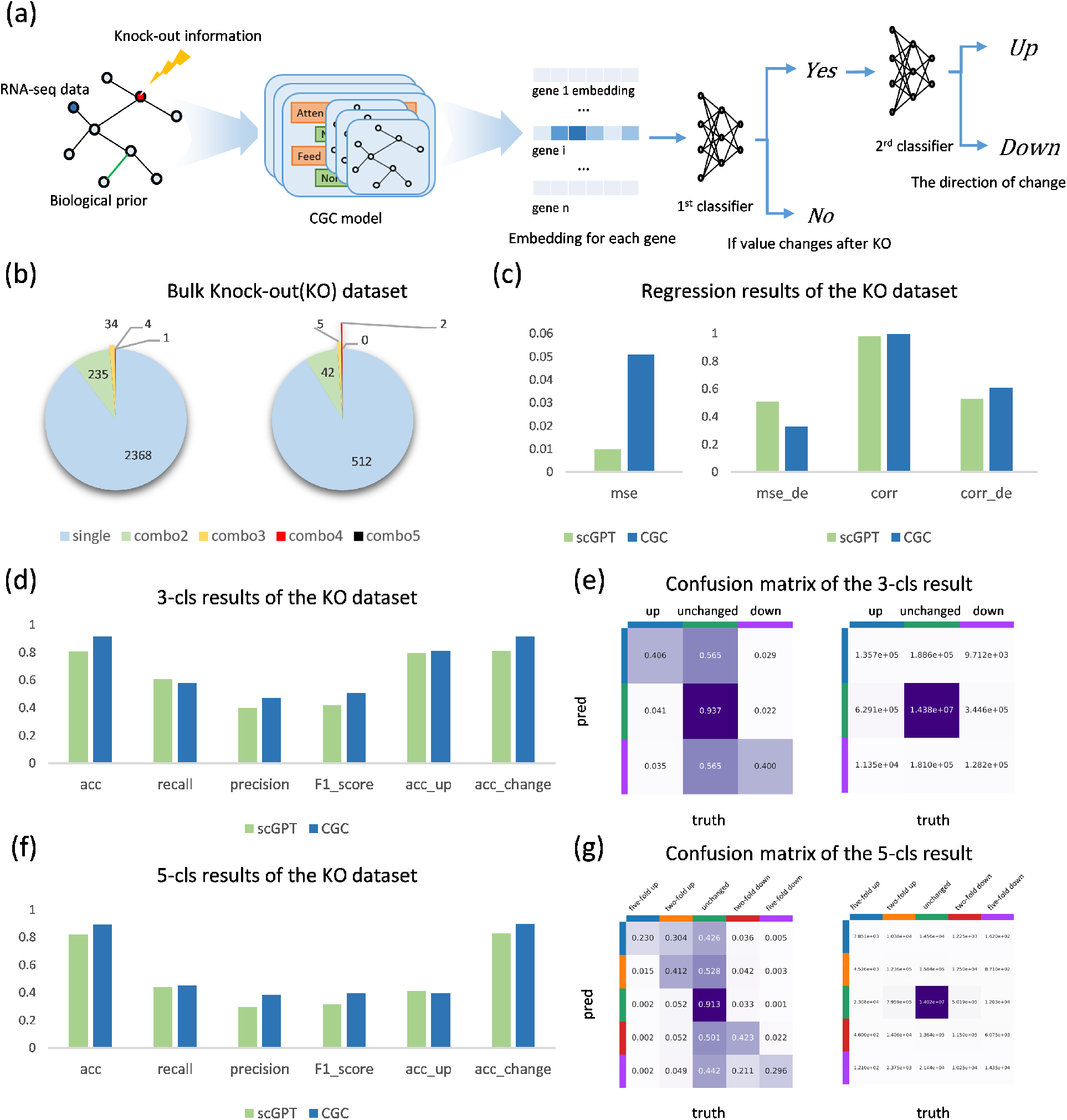
Exploring universal bulk gene knockout prediction with CGC. (a) Model architecture: We utilize a ”two-step” classifier as the decoder module of the CGC model for the bulk knock-out problem scenario. For each gene’s embedding output by the Transformer module, CGC first determines whether its expression value changes, then judges whether it is up or down. (b) The distribution of perturbation types in the bulk knockout dataset we collected. (c) Test results for the regression task. (d) Test results for the three-category (up, no change, down) task. (e) Confusion matrix for the three-category results, with the normalized on the left and the absolute values on the right. (f) Test results for the five-category (five-fold up, two-fold up, no change, two-fold down, five-fold down) task. (g) Confusion matrix for the five-category results.

In summary, CGC effectively transfers knowledge acquired from single-cell data pre-training to the bulk level, enhancing the performance of bulk gene knockout predictions, and demonstrates significant potential in guiding real-world biological research.

## 3 Discussion

In this article, we introduce Cell-Graph Compass, the pioneering single-cell domain foundation model utilizing graph structures to model genes and cells. By embedding various biological features and prior knowledge through Graph Neural Networks, we achieved results surpassing the state-of-the-art (SOTA) methods across a diverse range of downstream tasks. Our analysis of the pre-trained model’s latent space, including extensive gene classification and Gene Regulatory Network inference experiments, confirms that our pre-training captured an understanding of genes and their interactions to a certain extent. Cell clustering experiments demonstrate CGC’s effectiveness in overcoming batch effects, while cell type annotation tasks further validate CGC’s applicability to cell-level problems. Furthermore, we delved into gene perturbation response prediction, where CGC improved predictions for single-cell gene perturbations and showed potential in building a universal bulk-level gene knock-out prediction model, supported by a collection of 300,000 bulk RNA-seq control and over 3,000 bulk knock-out RNA-seq datasets for future research.

Through rigorous comparative and ablation studies, we demonstrated the efficacy of the CGC methodology. Comparisons with smaller models trained from scratch, along with ablation studies on the pre-training stage, underscored the effectiveness of pre-training operations. Evaluating against Transformer-only foundation models and ablation of the GNN module confirmed our approach to graph modeling. A wealth of experimental results demonstrate that CGC can be applied across a broad range of problem scenarios, achieving the best results to date.

Regarding the limitations of our work and the future directions, we present several considerations. Firstly, the information provided by transcriptomics alone is limited; we aim to evolve our single-cell RNA-seq foundation model into a comprehensive multi-omics integrated model, integrating data from ATAC-seq, proteomics, epigenomics and so on. Secondly, the features embedded in CGC can be further complexified and diversified. For example, currently utilizing gene positional information on chromatin as a one-dimensional edge feature is relatively simplistic; we plan to replace it with spatial transcriptomics data, which could offer richer structural information between genes. Moreover, with the rapid advancement of Large Language Models (LLMs),^66–68^ exploring more advanced LLMs beyond BioBERT for extracting textual information from gene descriptions is promising. Thirdly, our model architecture, which chains GNNs with Transformers, is relatively basic. Future work could experiment with alternating GNN and Transformer layers^69^ or other more complex and advanced architectures.^70, 71^ We might even explore constructing dynamic graphs for genes to incorporate temporal information. Lastly, to address the issues of data scarcity and imbalance met in downstream tasks, beyond using pre-training and rich biological features, semi-supervised learning^72–74^ and long-tail learning^75, 76^ may offer more solutions. In summary, CGC offers a methodology for constructing graph-structured foundation models using RNA sequencing data and biological features, indicating many directions and details worthy of further exploration.

As the ”pre-training & fine-tuning” paradigm gains momentum in life sciences, graph-based foundation models are poised to shine in areas like cell fate reprogramming, drug development for cancer treatment, organoid culture and so on, truly aiding wet-lab biology and becoming a staple in bioinformatics analysis.

## 4 Methods

### 4.1 Diverse and Comprehensive biological Features Used by CGC

Cell-Graph Compass effectively utilizes Graph Neural Network architectures for the integration of various feature types. Our encoding method distinguishes these features into two categories: node features and edge features. For each input RNA-seq data, CGC creates a distinct graph where nodes symbolize all the genes present in this cell, and edges represent the interconnections between these genes. Therefore, any information pertaining to individual genes is encoded as node features, while details depicting the relationships among genes are encoded as edge features.

### Node features

- Gene tokens: Each RNA-seq sample fed into CGC contains a combination of genes. To identify the unique identities of these genes, we have assigned a global ID to every gene encountered during the pre-training process. Analogous to the concept of word tokens in NLP, we refer to these gene IDs as ‘gene tokens.’ The use of gene tokens enhances the flexibility of the model’s input. For new genes, simply assigning a new ID allows their seamless integration into the model’s training framework. For existing genes, the iterative feature learning of gene tokens maintains the model’s ‘memory’ of these genes, ensuring continuity and precision in gene-related data processing.
- Gene Text Descriptions: Currently, there is an abundance of research focused on gene functions accompanied by extensive textual descriptions. Mining these existing textual resources is advantageous for rapidly acquiring some preliminary, foundational information about genes. We have obtained textual descriptions of various genes from the NCBI^8^ gene database, and these descriptions have been transformed into 768-dimensional feature vectors by querying BioBERT,^25^ which then serve as inputs for CGC.
- Gene Expression values: The expression levels of genes within a cell reveal the cell’s type and its current state. Providing the model with information on gene transcription expression is crucial for acquiring features that are relevant to the gene’s context. We have sourced raw readouts of gene transcription expression from various datasets, which then undergo a consistent process of quality filtering, normalization, and log transformation. In certain specific scenarios, which will be discussed later, additional steps such as selection of highly variable genes or discretization are also performed.
- Special Tokens:This feature is designed to facilitate the embedding of additional information in specific scenarios. For instance, in the context of perturbation prediction, we use a ‘0/1’ indicator to denote whether a gene has been perturbed or not. In this paper, we employ this feature exclusively in two downstream tasks: single-cell gene perturbation prediction and bulk knockout prediction.

Therefore, for each sample input into CGC, the node feature of the i-th gene can be represented as (*g*_*i*_, *t*_*i*_, *v*_*i*_, *s*_*i*_) , where i ranges from 0 to N-1. Here, N represents the number of genes contained in this sample. *g*_*i*_ ∈ ℕ denotes the global ID of the i-th gene, *t*_*i*_ ∈ ℝ^768^ represents the textual embedding of the i-th gene in 768 dimensions, *v*_*i*_ ∈ ℝ indicates the transcriptional expression of the i-th gene, and *s*_*i*_ stands for any potential special tokens.

### Edges features

- Transcription Factor-Target Gene Interactions:Research on transcription factors and their target genes has been conducted for many years. Embedding the results of previous research as prior knowledge into our model is expected to significantly enhance the performance of CGC. We have obtained pre-existing TF-target gene relationships from two industry-recognized databases: the TFTG database^77^ and the TRRUST database.^78^ The former contains TF-target gene regulatory relationships inferred from experiments, while the latter summarizes regulatory relationships documented in the literature. Based on this information, we generate a directed graph for each input sample, where the edges start from transcription factors and point to their target genes.
- Gene Co-expression Relationships:The activation patterns of genes in different cells reflect the functions and roles of these genes. If two genes exhibit highly correlated transcriptional expression profiles across a range of cells, it’s likely that they share similar functions or regulatory relationships. We employ the Pearson Correlation Coefficient (PCC) as a measure of gene co-expression relationships and retain gene pairs with a PCC greater than 0.6 to form the edges in the second graph.
- Positional relationship on chromosomes:Genes located in closer proximity on the same chromosome tend to exhibit higher similarity. We have calculated the relative distances between all gene pairs on the chromosomes (measured in terms of the number of genes separating them), and retained those gene pairs with a relative distance of less than 50 to form the edges in the third graph.

For each input sample, we create the aforementioned three subgraphs based on the genes it contains, which are then assembled to form a final, larger graph. The features of each edge on this graph can be marked as (*tftg*_*i*,*j*_, *corr*_*i*,*j*_, *pos*_*i*,*j*_), where the subscipt (*i*.*j*) represents the edge connecting the i-th node and the j-th node. Here, *tftg*_*i*,*j*_ ∈ {0, 1} indicates whether this gene pair has a TF-TG relationship, *corr*_*i*,*j*_ ∈ [0, 1] represents the PCC coefficient of the gene pair, and *pos*_*i*,*j*_ ∈ {−50, −49, …, 50} signifies the relative position of the gene pair on the same chromosome (with *pos*_*i*,*j*_ = 0 indicating that the relative position of the gene pair exceeds 50 or they are not on the same chromosome).

### 4.2 CGC model architecture

#### Embedding module

For the different types of feature data inputted into CGC, we have designed specific encoders to integrate all information into unified feature vectors for nodes and edges. Gene Tokens, Special Tokens, TF-TG interactions, and Positional relationship on chromosomes are encoded using traditional embedding layers *emb*_*g*_(·), *emb*_*s*_(·), *emb*_*tftg*_(·), *emb*_*pos*_(·) while Gene Text Descriptions, Gene Expression Values, and Gene Co-expression relationships are encoded using three-layers Multi-Layer Perceptrons(MLPs) *mlp*_*t*_(·), *mlp*_*v*_(·), *mlp*_*corr*_(·). The outcomes of these distinct encoding processes are then cumulatively combined to form the final comprehensive embedding representations for both nodes and edges, denoted as

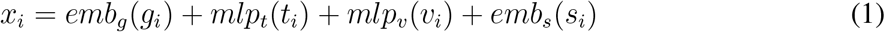

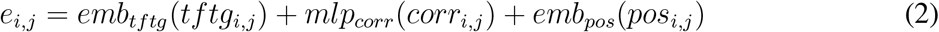

respectively.

#### GNN module

Adhering to the message passing paradigm, we have developed a three-layer Graph Neural Network (GNN) module to jointly learn the feature representations of nodes and edges. In each layer, the information of adjacent nodes and connecting edges is utilized to update the central node. Subsequently, the central edge’s information is updated using the data from the edge itself and its connected nodes. The process is articulated through the following formulas:

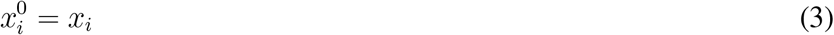

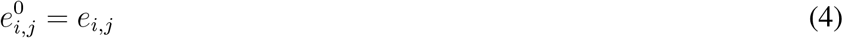

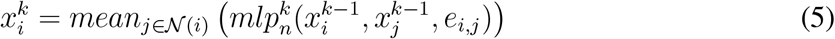

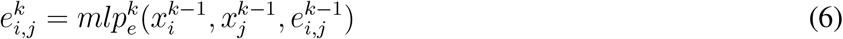

where *k* ∈ {1, 2, …, *L*} and *L* represent the number of GNN layers. 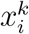 and 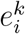 represent the representation vectors of nodes and edges output by the k-th layer, with 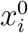 and 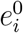 denote the input of the GNN module. 𝒩 (*i*) refers to the set of neighboring nodes of gene *i*.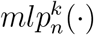 and 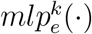 both refer to a three-layer Multi-Layer Perceptrons while *mean*(·) denotes the operation of calculating the average.

A single layer of graph convolution updates the target using information from adjacent nodes and edges and three concatenated layers of graph convolution leverage information from all nodes and edges within a three-hop range to update the target. The final output of the GNN module’s integrates local information related to the target node.

#### Transformer module

Next, we designed a self-attention Transformer module to capture the interactions between genes from a global perspective. Specifically, we first derive the query, key, and value matrices for the self-attention operation from the node features outputted by the GNN:

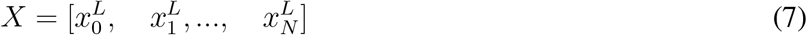

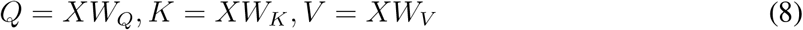

where *W*_*Q*_, *W*_*K*_, *W*_*V*_ ∈ ℝ ^*d′×d*^ are learnable parameters, with *d*^*′*^ and *d* is the dimension of the hidden vectors. Then, we apply the self-attention mechanism to model the interrelationships among all nodes:

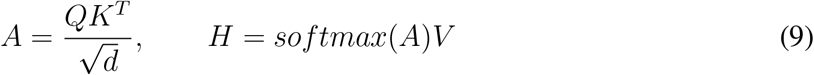

Here, *H* = [*h*_0_, *h*_1_, …, *h*_*N−*1_] represents the feature representation outputted by the transformer module for each gene.

In addition to the traditional transformer model architecture, we have also implemented Flash-Attention^79^ technology to accelerate the attention mechanism. The Flash-Attention technique significantly reduces training costs, enabling the Transformer to handle inputs consisting of thousands or even tens of thousands of tokens.

#### Cell embedding

Once we have obtained the representations for all the genes within a sequencing sample, we can synthesize these to form an overall representation of this cell. This could be achieved by simply averaging or performing a weighted sum of all genes’ feature vectors. However, in this paper, we adopted a more sophisticated approach. We introduced an additional node to the graph representing the overall state of the cell, with all other nodes representing genes connected to it. This allows both the GNN and the Transformer module to autonomously learn the integrated representation from the gene representations. The feature representation *h*_*c*_ outputted by the Transformer for this node is then extracted as the cell embedding.

### 4.3 The pre-training of CGC

#### Assembling and preprocessing pretraining corpus

To support foundation model research, we have constructed a large-scale single-cell transcriptome corpus, ScCompass-126M.^16^ It contains 126 million single-cell transcript sequencing entries sourced from various tissues and organs of humans and mice. In this study, we utilized about 50 million of these entries from human samples to conduct our pre-training. For the purpose of quality control, we excluded cells that were either of low quality or damaged and then all entries were standardized and log-transformed to ensure consistency and accuracy. To enhance the efficiency of our pre-training, we initially selected 10,000 highly variable genes for each SRR experiment separately. Subsequently, for each sequencing entry, we sampled 1,200 genes based on their expression values to serve as inputs for the pre-training model. Additionally, in order to mitigate batch effect, during the pre-training stage, we employ the value binning technique proposed in scGPT^10^ to convert all gene expression values into discrete relative values ranging from 0 to 50.

#### CGC pre-training strategy

We employed a traditional masked training strategy for self-supervised learning from a large-scale, unlabeled dataset. For each input sample, we randomly mask 40% of the gene expression values and predict these masked values using the expression of the remaining genes. This process can be formalized as follows:

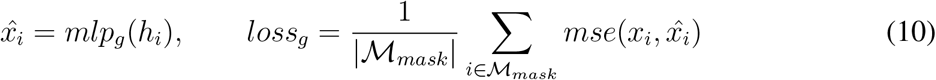

where *x*_*i*_ and 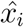 represent the actual and predicted expression values of the i-th gene in each sample, and *h*_*i*_ is the hidden vector output by the Transformer module. ℳ_*mask*_ is the set of indices for the masked genes, and *mse*(·) denotes the mean squared error operation.

For the learning of cell embedding, we attempted to predict the expression values of all genes using cell embedding and gene tokens. The process is formalized as follows:

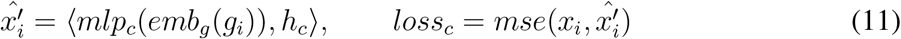

where *h*_*c*_ represents the cell embedding , *emb*_*g*_(*g*_*i*_) denotes the gene token embedding, and ⟨·, ·⟩ signifies the inner product operation.

### 4.4 Downstream tasks

#### Gene Classification

To explore whether CGC’s pre-training can learn biological knowledge, we extracted the gene token encoder’s embeddings, *emb*_*g*_(·), for each gene. Initially, these embeddings are randomly initialized, but after pre-training, they should contain a global understanding of these genes. We tested the ability of CGC’s gene embeddings to identify the identity or characteristics of specific genes using six gene classification datasets compiled by Theodoris et al.^9^

- Dosage sensitive prediction tests whether a gene is sensitive to changes in its expression level, crucial for understanding the mechanisms of genetic diseases caused by gene dosage imbalances. We used previously reported^17–19^481 genes for experiments.
- Bivalent chromatin structure prediction involves a unique epigenetic modification at some gene promoters, significant in gene expression regulation, especially during early development and stem cell differentiation. The bivalent domain combines two opposing histone modifications: H3K4me3 and H3K27me3. Using data on 56 conserved regions from previous reports,^20^ we first differentiated between bivalent and non-methylated genes, then further between bivalent and Lys4-only methylated genes, with dataset sizes of 147 and 187, respectively.
- Transcription factors’ action range prediction reflects different regulatory modes by the distance of TF binding to target DNA sequences. We utilized a dataset from Chen et al.,^21^ comprising 173 samples.
- Network dynamics prediction involves identifying core genes within gene regulatory networks, a key to understanding the mechanisms of certain diseases. We utilized a NOTCH1 (N1)-dependent gene network provided by Theodoris et al.,^22, 23^ aiming to predict which genes are core to the N1 network and which are peripheral downstream effectors. Furthermore, we sought to predict which genes are influenced by this N1 network. The sizes of the two datasets are 281 and 1103, respectively.

We generated embeddings for each gene using Gene2vec, BioBERT, and CGC, along with random embeddings for comparative experiments. For each model’s embedding, we uniformly employed a random forest classifier, evaluating performance through five-fold cross-validation on the afore-mentioned datasets.

ROC and AUC served as testing metrics. The Receiver Operating Characteristic (ROC) curve graphically shows the relationship between True Positive Rate (TPR) and False Positive Rate (FPR) at various thresholds. The Area Under the Curve (AUC) quantifies the performance of classification models. With AUC values ranging from 0 to 1, a perfect model scores 1, indicating flawless distinction between classes at all thresholds. An AUC value of 0.5 suggests performance equivalent to random guessing. Generally, closer AUC values to 1 indicate better model performance.

#### Gene Regulatory Network Inference

The GRN inference task aims to predict the interactions between transcription factors and target genes directly. Our data is derived from two databases, STRING^28^ and ChIP-seq,^29–31^covering two cell types: human embryonic stem cells (hESC) and human mature hepatocytes (hHEP). Following work by Pratapa et al.,^32^ we experiment with the 500 and 1000 genes respectively, which show the highest variance among all TFs with a corrected P-value of variance below 0.01. Similar to the inference method of DeepSEM,^33^ we use the cosine similarity between gene pairs as the basis for prediction:

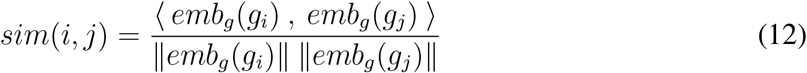

where ⟨·, ·⟩ signifies the inner product operation and ∥·∥ represents the calculation of the Euclidean norm. We evaluate the model’s ability to reconstruct the GRN in two ways. First, we retain the top *K* gene pairs with the highest similarity, where *K* represents the number of edges in the ground-truth network. We then calculate the proportion of our retained *K* gene pairs that overlap with the edges in the ground-truth network, as depicted by the Early Precision Ratio (EPR) metric. Secondly, based on the similarity of gene pairs, we have the model output the probability that there is a regulatory relationship between them, quantified by the Area Under the Precision–Recall Curve (AUPRC) ratio, to measure the model’s capability to predict edges present in the ground-truth network.

The extraction of gene programs was conducted on the human immune tissues dataset. Our validation steps follow the approach of Cui et al.^10^ : Firstly, we selected highly variable genes and constructed a gene similarity network based on the similarity between gene pairs, as with Equation Subsequently, we applied Louvain clustering^80^ to the resulted network at a resolution of 40 in order to divide these genes into different gene programs. Finally, we calculated the average expression of each program across various cell types.

#### Cell clustering and batch effect correction

The task of cell clustering aims to differentiate between data from various cell types across different sequencing batches. We employed the same data preprocessing procedure and masked training method as in the pre-training phase. In addition to the expression profile reconstruction loss mentioned in Equation 10 and 11, we also utilized the Domain Adaptation loss via Reverse Back-Propagation and a loss related to Elastic Cell Similarity, as proposed in scGPT^10^ . For comparison, we chose scVI, scFoundation and scGPT against CGC, with quantitative comparison standard derived from the biological conservation and batch effect correction metrics proposed by Luecken et al.^34^ :

- Normalized Mutual Information (NMI):NMI is used to measure the amount of shared information between clustering results and cell type labels. Through normalization, the value of NMI is constrained between 0 and 1, where 0 indicates that the two datasets are completely independent, and 1 indicates that the datasets are completely related or the information is fully shared.
- Adjusted Rand Index (ARI):The Rand Index (RI) is an indicator used to measure the consistency between clustering results and cell type labels by comparing the consistency and inconsistency of paired combinations in the clustering allocation. ARI is an improved version of RI, adjusted to account for the effects of random classification. The value of ARI ranges from 0 to 1, where 1 represents a perfect clustering effect, and 0 indicates the result of random clustering.
- Average Silhouette Width(ASW):The silhouette coefficient is used to calculate the difference between the average distance of a sample to other samples in the same category (cohesion) and the average distance to samples in the nearest other category (separation). We use batches as category labels and take the complement of the generally defined ASW as our indicator for depicting batch confusion: *ASW* = 1 − |*ASW*_*batch*_|. Under this definition, the ASW ranges from 0 to 1, with higher values representing better batch effect correction.
- Graph Connectivity (GraphConn):This metric measures the average proportion of samples within each cell type that are connected through the K-Nearest Neighbors (KNN) method. The value of Graph Connectivity ranges from 0 to 1, representing the model’s ability to overcome batch effects.

The average of these four metrics is considered the overall score.

#### Cell type annotation

Cell type annotation is a classification task at the cellular level. Our objective is to predict each sample’s cell type based on its gene expression profile. CGC employs a three-layer MLP as the decoder for this scenario, which uses the cell embeddings *h*_*c*_ as input to generate predictions. This process can be mathematically represented as follows:

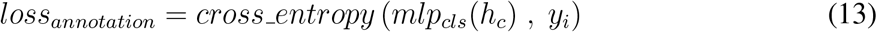

where *y*_*i*_ and *mlp*_*cls*_(*h*_*c*_) respectively represent the actual and predicted values of this sample’s cell type where *cross entropy*(·) represents the operation of calculating cross-entropy. We employ traditional classification metrics such as *accuracy, precision, recall*, and *F* 1 *score* to quantitatively evaluate the performance of tested models. Notably, the *accuracy* metric is calculated globally (i.e., micro), while the other metrics represent the average outcomes across different cell types (i.e., macro).

#### Single-cell gene perturbation prediction

The perturbation prediction task can be defined as follows: given a set of cells’ gene expression profiles in control condition (unperturbed) and perturbation conditions (which genes will be disturbed), predict their respective gene expression profiles post-perturbation. In this application scenario, we employ the special tokens mentioned in Method 4.1 and adopt the complete transcript expression values as input. We add a new one-hot encoded feature to each node, indicating whether the gene was directly perturbed in the experiment. CGC outputs an embedding representation for each gene and uses a shared decoder across genes to determine their post-perturbation expression levels. During the fine-tuning process, the loss function for each sample is defined as

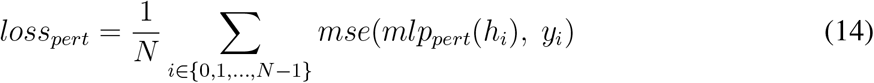

where N is the number of genes this sample contains, *mlp*_*pert*_ denotes the decoder and *y*_*i*_ represents the ground-truth value of post-perturbation expression.

To quantify the model’s predictive accuracy, we calculated the mean squared error (mse) and pearson correlation coefficient (corr) between the actual and predicted values of all genes in the test set. Since most genes show subtle changes pre- and post-perturbation, we specifically considered the top 20 genes with the most significant changes (DE genes), calculating their corresponding mse and corr, denoted as mse de and corr de. Furthermore, to amplify the differences between different tested models, we calculated the corr between the actual and predicted changes in gene expression, termed corr delta.

Note that the above test results are based on the average outcomes for the entire test set. Due to the nature of perturb-seq technology, in a single perturbation experiment, the gene expression profiles under the same perturbation condition are very similar. Therefore, we further calculated test results based on the average per perturbation condition. Specifically, we categorized the samples in the test set according to perturbation conditions, then calculated the mse, corr, and corr delta between the actual and predicted values for each perturbation condition, before averaging these metrics across all perturbation conditions. This testing method continues the approach used in GEARS.

#### Bulk-level gene knockout prediction

The bulk-level gene knockout is a downstream task that, while similar in definition to single-cell gene perturbation, holds an entirely different significance. In single-cell perturbation experiments, our focus is often limited to samples under specific cell conditions, with an emphasis on enhancing prediction accuracy. However, in bulk knockout prediction experiments, our aim is to develop a more universal predictive model applicable to a broader range of cell conditions, in hopes of uncovering universal relationships between genes.

To transfer the single-cell pre-trained model to the bulk knockout scenario, we employed a strategy of secondary pre-training. We downloaded 700,000 high-throughput sequencing data entries of mice from the ARCHS4 database,^81^ retaining 300k after quality filtering to serve as our bulk-level pre-training dataset. These sequencing data are organized in FPKM format and have undergone log1p transformation and sampling based on expression values, among other processes. The final input for the model consists of 2048 gene tokens per entry.

In building the bulk knockout dataset, we initially extracted sequencing results related to knock-out experiments from NCBI’s GEO database.^8^ We then categorized these sequencing results individually based on the experimental condition according to overall design of these GSE experiments and the different treatments of GEM samples. Ultimately, we retained the average expression of all samples under each experimental condition as a single entry in our bulk knockout dataset. Hence, some entries in our dataset, despite having the same knocked-out gene, exhibit significant data distribution differences due to their varying experimental conditions. Approaches like GEARS, which rely solely on the identity of the knocked-out gene for prediction, are clearly inapplicable in this scenario.

We first retrained our single-cell pre-trained model on the 300k bulk sequencing data using the same masked training loss mentioned in the Formula 10 and 11, and then fine-tuned it on the bulk knockout dataset using Formula 14. Although the experimental results surpassed those of comparative state-of-the-art (SOTA) models, we couldn’t discern their practical applicability. Therefore, we further shifted our research question to directly predict the direction of expression changes following gene knockouts, fine-tuning the model with classifiers and cross-entropy loss function. To address the issue of too few ’up-/down-regulated genes in the data, we adopted a ”secondary prediction” strategy. For each gene, we first predict whether it will change, and then for each gene that changes, we predict the direction of its change. This process is expressed in the following formula:

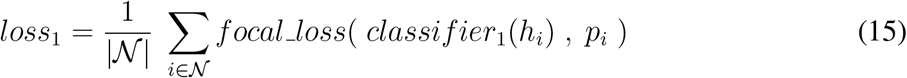

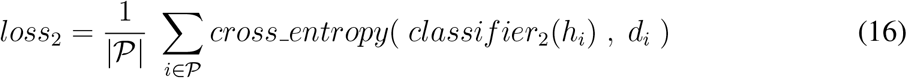

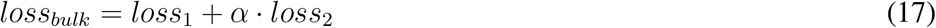

where 𝒩 represents the set of genes in this sample, and 𝒫 denotes the set of changed genes after knockout. *α* is the weight coefficient that coordinates the proportion between *loss*_1_ and *loss*_2_. *p*_*i*_ indicating whether the gene changes, and *d*_*i*_ specifying the direction of the change. *focal loss* is a specialized form of weighted cross-entropy. We utilized traditional classification metrics for quantitative comparisons and presented our predictive capability for changing genes using confusion matrices.

### 4.5 Implementation Details

The GNN module of the pre-training model is stacked with 3 layers, and the Transformer module is stacked with 12 blocks with 8 attention heads each. The embedding size and the fully connected layer hidden size are both 512. We use scCompass-h50M^16^ dataset for pre-training, where 99% of the data is used for training and 1% for validation. We use the mini-batch size of 32 with the gradient accumulation step number for 2, and a total of 8 gpus were used for 3 epochs of training. Note that only non-zero expression genes are input in the pre-training stage, and the max sequence length was set to 1200. The mask ratio is set to 0.4 and the ratio of *loss*_*g*_ (eq10) to *loss*_*c*_ (eq11) is set to 1. The model uses the Adam optimizer with Adam betas set to (0.9,0.999) and eps of 1e-8. The model used warmup and exponential decay to adjust the learning rate. The max learning rate is set to 1e-4, the warmup steps is 5000, the decay steps is 1000, and the decay ratio is 0.995.

For the fine-tuning downstream tasks, we keep the same model configuration with the pre-trained model. The cell clustering task uses the batch size of 32 with gradient-free accumulation, and the max sequence length of 1200. The ratio of the training set to validation set is 9:1, and the the ecs threshold of 0.8. For the cell type annotation task, the batch size is set to 8 with the gradient accumulation step number for 2 and the max sequence length is set to 3000. The training set and the validation set were divided by 9:1 on the reference dataset, and 30 epochs of training were conducted using 2 gpus. For single-cell gene perturbation prediction task, the batch size is set for 64 with gradient-free accumulation, and 30 epochs of training were conducted using 1 gpu. As for the bulk-level gene knockout prediction task, the focal loss weight is set to 1, focal alpha is set to 0.5, and other parameters are the same as those of single-cell gene perturbation prediction task.

Both pre-training and fine-tuning experiments used the machine with A100 40G gpu, 380G of machine memory, and 16 CPU cores, and the main code used the Uni-core framework. Scanpy and anndata are applied to pre-process single-cell data. And the LMDB package is used to write pre-processed data into memory for accelerating data reading. We use pyg to implement message passing mechanism and graph data processing, and flash attention^79^ to avoid length limitation of Transformer. The metrics of cell clustering are calculated and shown by scib.metrics,^34^ and the metrics of cell type annotation tasks are calculated by scikit-learn and shown by seaborn. The analysis of single-cell gene perturbation prediction and bulk-level gene knockout prediction is implemented by the cell-GEARS^60^ package.

### Disclosures

The authors declare no competing interests

## Code, Data, and Materials Availability

The code and data involved in this study will be made publicly available on GitHub once they are ready, and updates will be synchronized to this preprint.

## Acknowledgments

This work is supported by the Pilot Project for Enhancing Original Innovation Capability of the Chinese Academy of Sciences (E129L111) and Informatization Plan of Chinese Academy of Sciences (CAS-WX2021SF-0101)

## Supplementary Material

**Fig S1:**
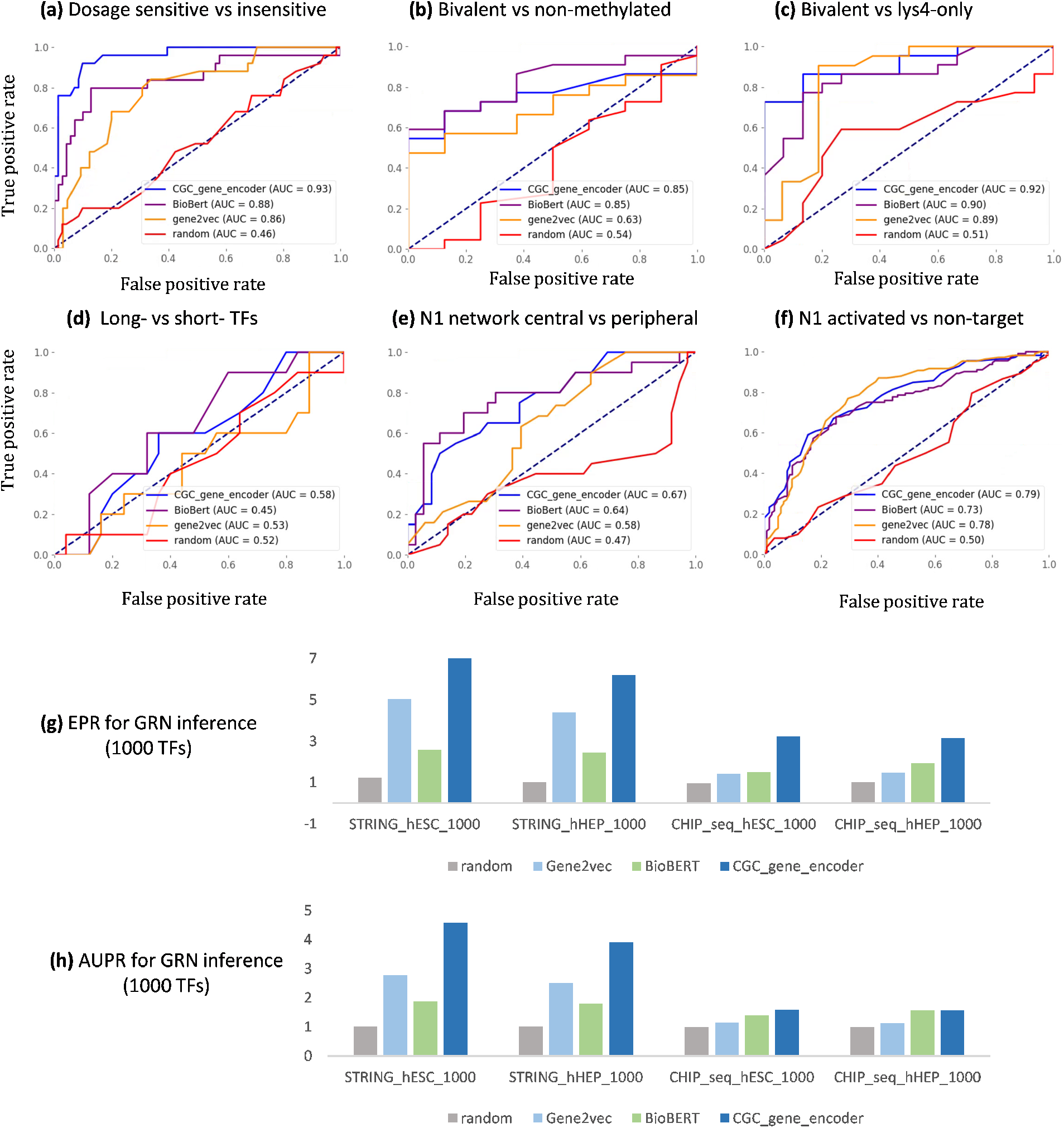
(a) to (f) ROC curves of the CGC Encoder and baseline models on six gene classification tasks. (g) & (h) Results of the CGC Encoder and baseline models in the inference experiments of GRNs for 1000 TFs, tested from the aspects of EPR and AUPR respectively.

**Fig S2:**
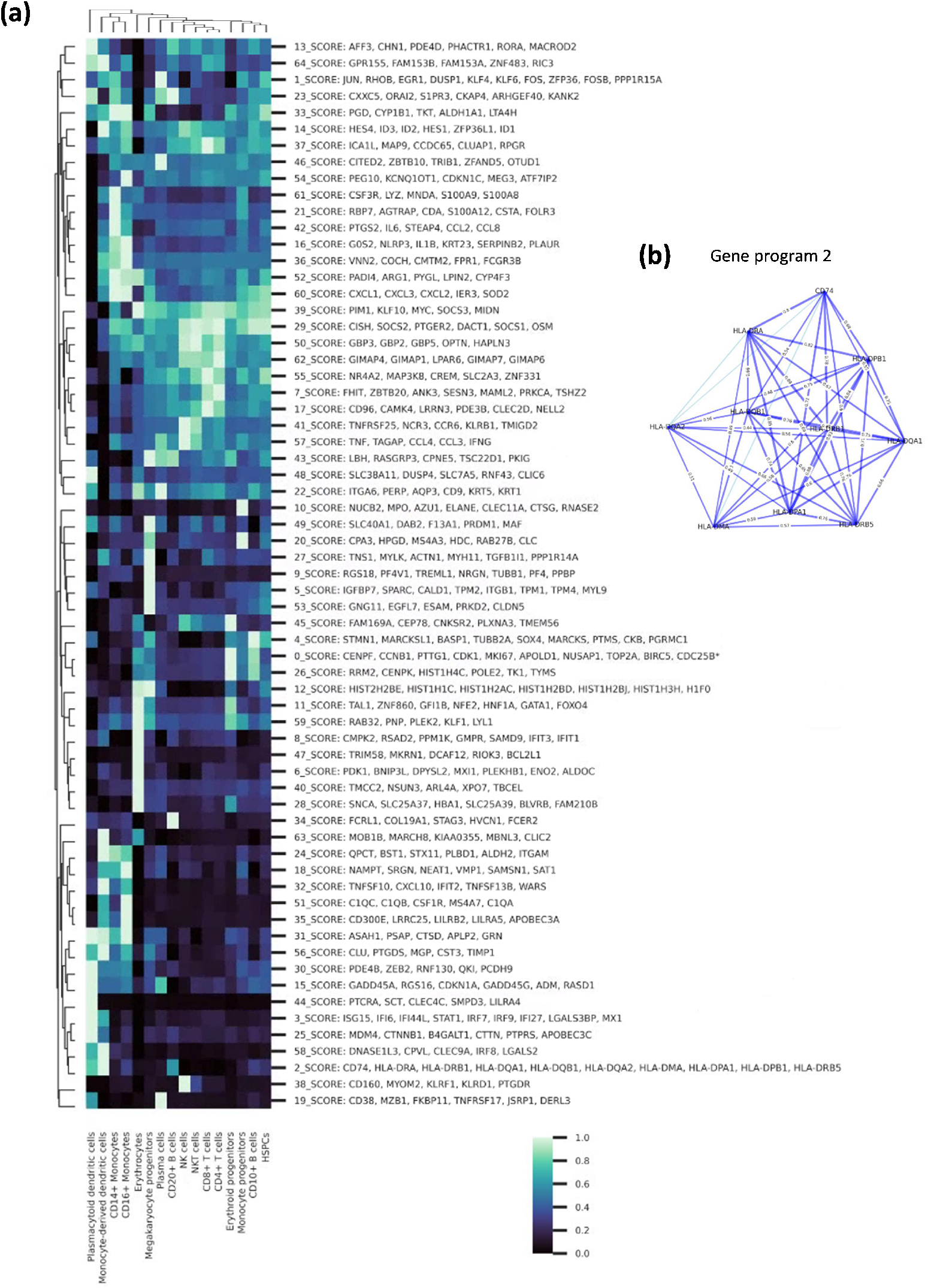
(a) Gene programs extracted by CGC and their selective expression across different cell types. (b) A visualization example: The gene regulatory network constructed by CGC within gene program 2.

**Fig S3:**
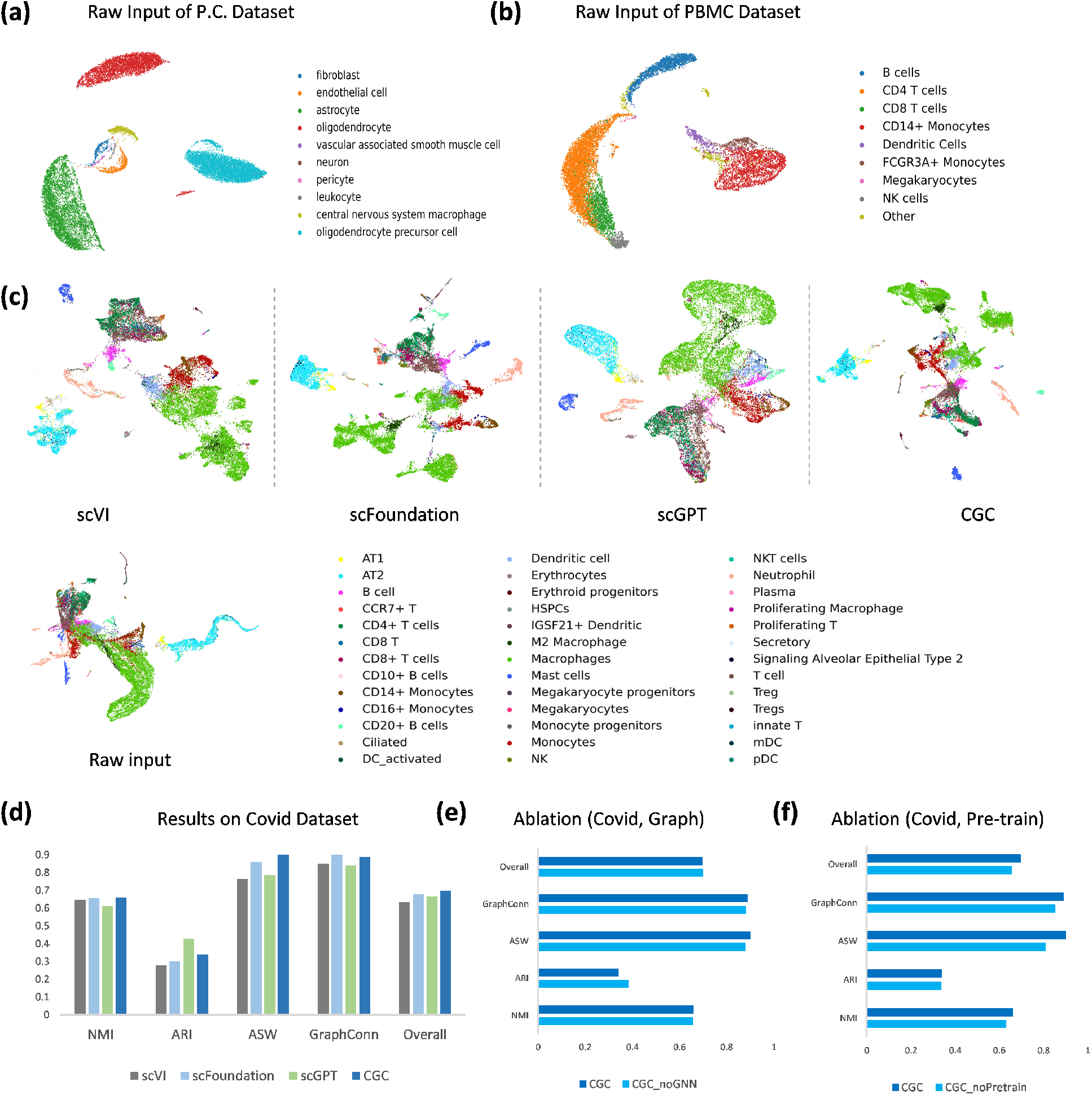
Supplementary figure for the Batch Effect Correction task. (a) UMAP plot of the original inputs for the PCortex dataset, colored by ground truth cell types. The original input refers to the vector composed of the expression values of all genes in the cell. (b) UMAP plot of the original inputs for the PBMC dataset. On the COVID dataset, (c) UMAP plots of the original inputs and the cell embeddings generated by all models. (d) Benchmark results for CGC and the baseline models. (e) Ablation study on the graph structure of CGC. (f) Ablation study on the pretraining process of CGC.

**Fig S4:**
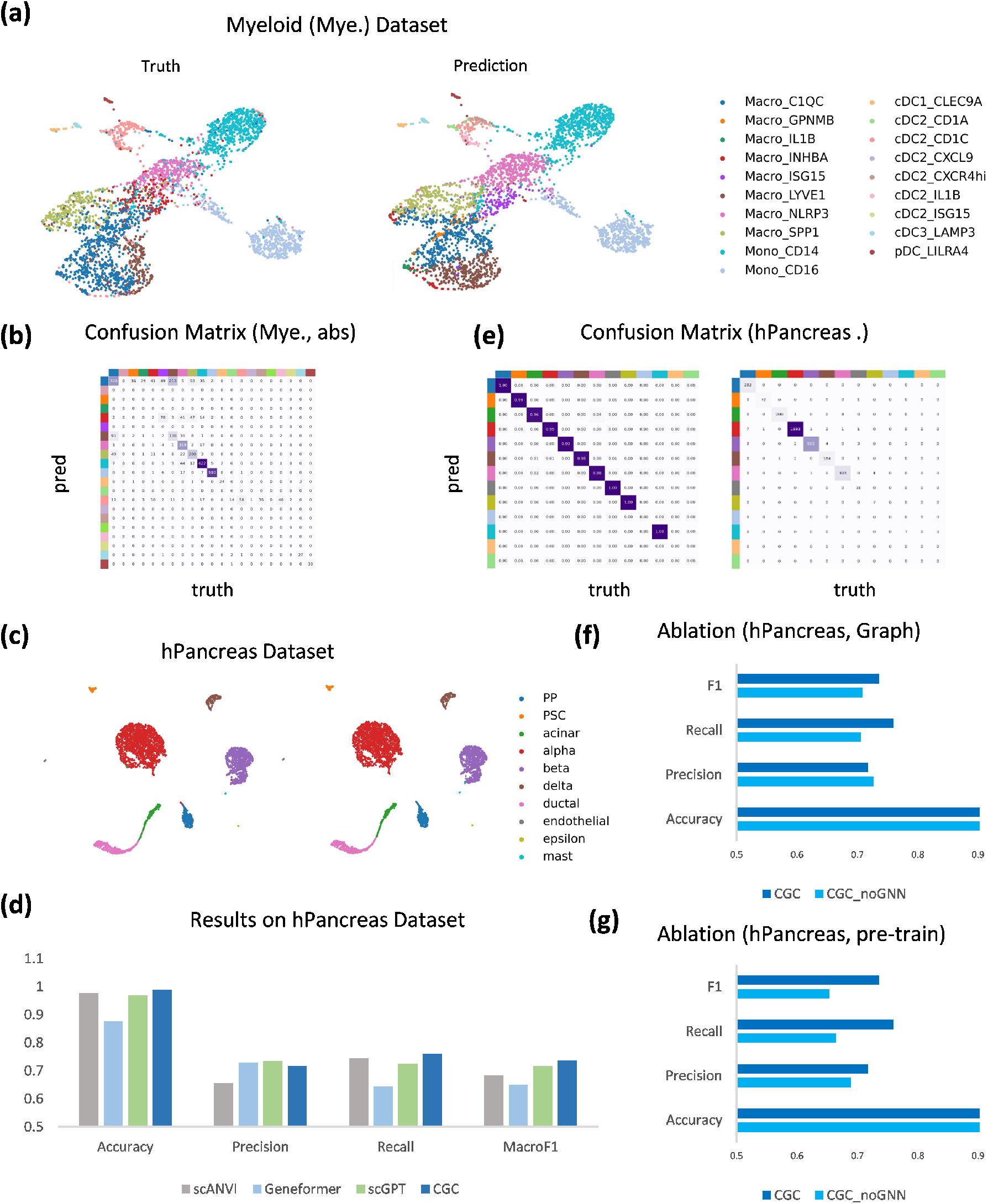
Supplementary figure for the Cell Annotation task. On the Myeloid (Mye.) dataset, (a) UMAP plot of cell embeddings generated by CGC, colored by ground truth cell types (left) and CGC prediction results (right). (b) Confusion matrix (absolute values) between CGC-predicted cell types and ground truth labels. On the hPancreas dataset, (c) UMAP plot of cell embeddings generated by CGC, colored by ground truth cell types (left) and CGC prediction results (right). (d) Quantitative evaluation of cell annotation by CGC and baseline models. (e) Confusion matrix between CGC-predicted cell types and ground truth labels, with normalization on the left and absolute values on the right. (f) Ablation study on the graph structure of CGC. (g) Ablation study on the pre-training process of CGC.

**Fig S5:**
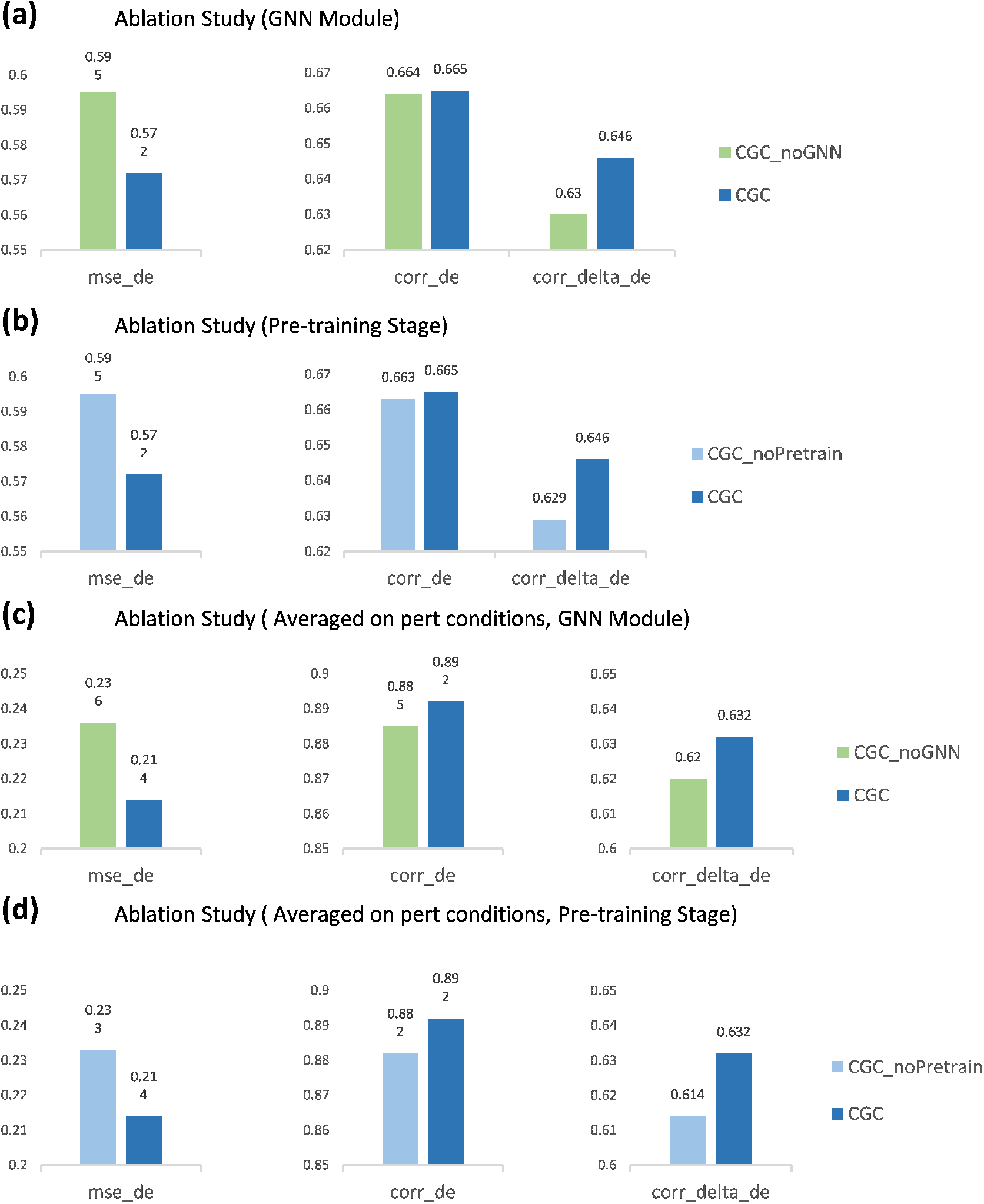
Ablation experiments of CGC on the task of single-cell gene perturbation response prediction. On the test set of the Norman dataset, (a) Ablation study on the graph structure of CGC. (b) Ablation study on the pre-training process of CGC. In experiments predicting the average response of genes under various perturbation conditions, (c) Ablation study on the graph structure of CGC. (d) Ablation study on the pre-training process of CGC.

**Fig S6:**
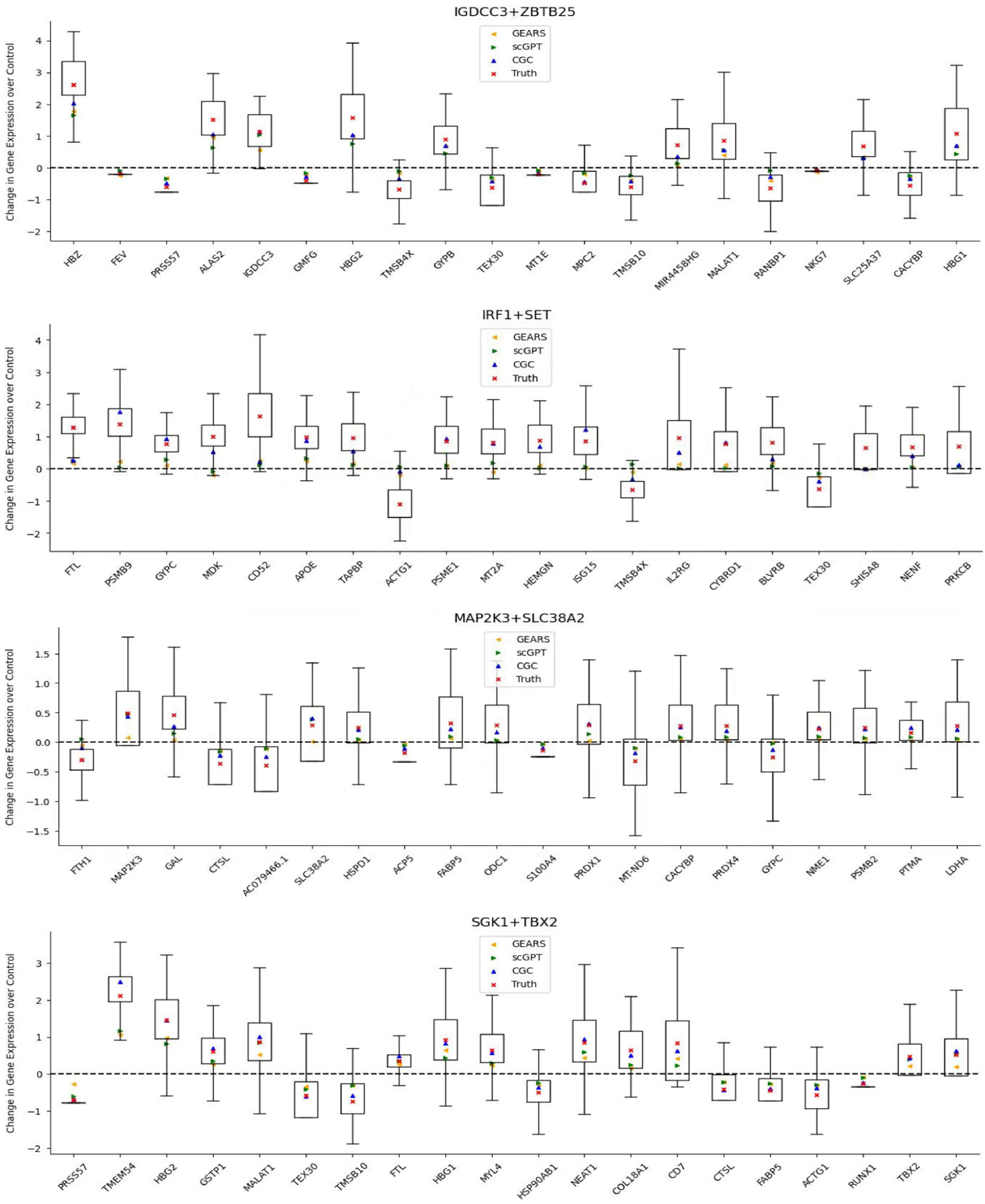
Examples of gene perturbation response prediction. From top to bottom, the perturbation conditions are for IGDCC3 and ZBTB25, IRF1 and SET, MAP2K3 and SLC38A2, SGK1 and TBX2, respectively. The x-axis shows the top 20 differentially expressed (DE) genes under each perturbation condition, and the y-axis indicates the change in gene expression due to the perturbation. The box plots represent the distribution of real experimental data for each perturbation condition, with red dots indicating their mean values. Other colored dots represent the prediction values from various models.

